# A Meta-Analysis of the Effects of Acute Sleep Deprivation on the Cortical Transcriptome in Rodent Models

**DOI:** 10.1101/2025.04.21.648791

**Authors:** Cosette A. Rhoads, Megan H. Hagenauer, Jinglin Xiong, Erin Hernandez, Duy Manh Nguyen, Annaka Saffron, Elizabeth Flandreau, Stanley Watson, Huda Akil

## Abstract

Sleep deprivation (SD) causes large disturbances in mood and cognition. The molecular basis for these effects can be explored using transcriptional profiling to quantify brain gene expression. In this report, we used a meta-analysis of public transcriptional profiling data to discover SD effects on gene expression that are consistent across studies and paradigms. To conduct the meta-analysis, we used pre-specified search terms related to rodent SD paradigms to identify relevant studies within *Gemma*, a database containing >19,000 re-analyzed microarray and RNA-Seq datasets. Eight studies met our systematic inclusion/exclusion criteria. These studies characterized the effect of 18 SD interventions on gene expression in the mouse cerebral cortex (collective *n*=293). For each gene with sufficient data (*n*=16,290), we fit a random effects meta-analysis model to the SD effect sizes (log(2) fold changes). Our meta-analysis revealed 182 differentially expressed genes in response to SD (false discovery rate: FDR<0.05), most of which (115/182) showed similar effects (FDR<0.05) in an independent large dataset (*GSE114845*: *n*=86 RNA-Seq samples from *n*=222 mice). Gene-set enrichment analysis revealed down-regulation in pathways related to stress response (e.g., glucocorticoid receptor *Nr3c1*), vasculature, growth and development, and upregulation related to stress, inflammation, and neuropeptide signalling. Exploratory analyses suggested that recovery sleep (included in six contrasts: range: 1-18 hrs), could reverse the impact of SD on gene expression. Our meta-analysis provides a useful reference database illustrating the diverse molecular impact of SD on the rodent cerebral cortex.

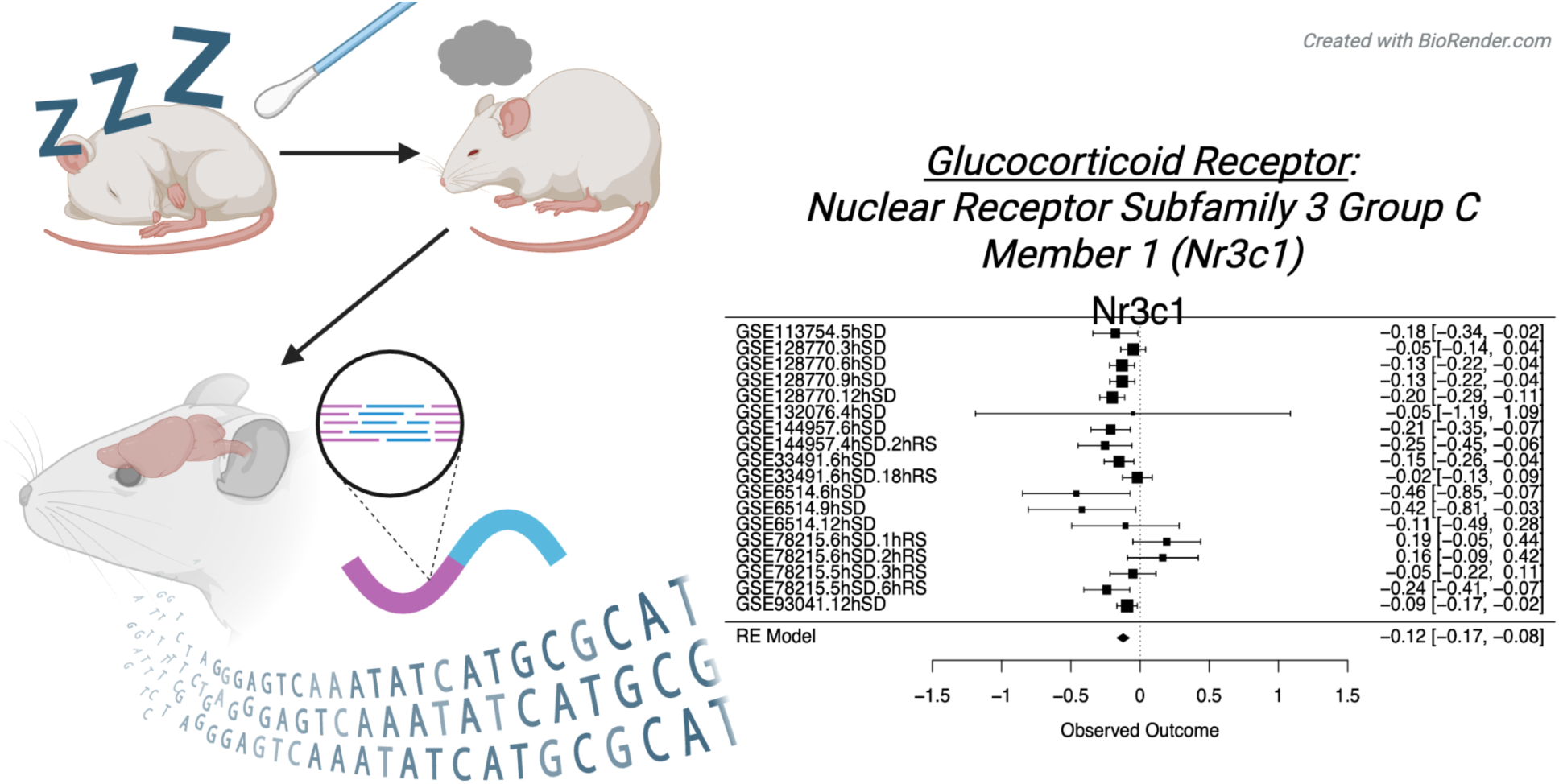
Graphical Abstract.

## Introduction

Sleep plays a critical role in many diverse physiological functions, including growth, neuroplasticity, protein synthesis, energy optimization, waste removal, repair and immune function (Elliott et al., 2014; Weiss and Donlea, 2021; Zielinski et al., 2016). Consequently, sleep disruption, restriction, or deprivation (SD) negatively impact many aspects of routine brain function, causing mood disturbances, impaired memory and cognitive abilities (Lyons et al., 2023; McEwen and Karatsoreos, 2022; Medic et al., 2017; Van Dongen et al., 2003). During SD, homeostatic drive increases the propensity to sleep. When sleep is eventually re-initiated, there is a compensatory rebound in the duration and intensity of sleep, especially during slow wave sleep (SWS) and rapid eye movement (REM) sleep stages, accompanied by a particularly large potentiation of cortical slow wave activity (SWA) (Borbély et al., 1981; Tobler and Borbély, 1990).

To gain insight into the molecular basis for these effects, gene expression can be quantified in the cortex following SD using transcriptional profiling methods that measure mRNA such as microarray and RNA-Seq. Transcriptional profiling provides rich information about cellular and tissue function but also presents many challenges, including methodological variability due to sample mislabeling (Toker et al., 2016), batch effects (Zhou et al., 2019), and dissection heterogeneity (Hagenauer et al., 2018). Additionally, the small sample sizes common in transcriptional profiling experiments provide low statistical power, increasing the risk of false negative and false positive findings (Button et al., 2013).

To overcome these challenges, we used meta-analysis to identify consistent effects of SD on the cerebral cortex within public transcriptional profiling datasets. Our meta-analysis focused on the effects of SD in laboratory rodents, induced by various methods for waking the animal soon after sleep initiation (“enforced wakefulness”). As rodents sleep in multiple short episodes throughout the day, rodent SD interventions tend to occur on the scale of hours instead of days (Pires et al., 2016), with longer periods of SD expected to produce more dramatic effects on the brain and behavior (Tobler and Borbély, 1990). To increase the generalizability of our findings, we included studies encompassing a variety of SD paradigms and subject characteristics in our meta-analysis. When possible, we used standardized, systematic methods for dataset identification, selection, and result extraction to reduce bias and improve the quality of our results. Finally, we validated our results using a separate large RNA-Seq dataset that was not included in our meta-analysis.

## Methods

### General Overview

The R code (R v.4.2.0, R studio v.2022.02.4) and detailed methods for our meta-analysis can be found at: https://github.com/rhoadsco/Sleep-Deprivation-MetaAnalysis. This meta-analysis was completed as part of the *Brain Data Alchemy Project*, a collective effort to improve the reliability and generalizability of transcriptional profiling findings using meta-analyses of public datasets. This guided effort uses a standardized pipeline for dataset identification, inclusion/exclusion, and meta-analysis (pipeline for 2022: (Hagenauer et al., 2024a), *not pre-registered*), leveraging the data curation, preprocessing, and analysis efforts of the *Gemma* database (Lim et al., 2021; Zoubarev et al., 2012). *Gemma* currently houses >19,000 curated and re-analyzed transcriptional profiling datasets, with an emphasis on brain-derived data.

We have provided a detailed description of Gemma’s data processing procedures in the supplementary methods for easy reference (*summarized from* (Lim et al., 2021a)). In brief, Gemma performs up-to-date RNA-Seq read mapping and microarray probe alignment to the reference genome. Alignments are filtered for specificity, and mapped to transcripts using UCSC Golden Path. Gene-level and sample-level quality control is conducted using standardized procedures applied similarly across platforms, including the removal of genes lacking variance in expression and outlier samples with low sample-sample correlations. Batch information from the metadata and raw data files is screened for potential impact using the correlation with top principal components of variation in the expression data. Batch correction is conducted using ComBat (Johnson et al., 2007), unless the batches confound the experimental design, in which case the dataset is flagged or split. Differential expression analyses are performed using the *limma* pipeline, followed by empirical Bayes correction (Ritchie et al., 2015). For RNA-Seq data, the voom algorithm is used to compute weights from the mean–variance relationship of the data (Law et al., 2014). Statistical output is available at the level of the omnibus model and individual group-level contrasts.

### Meta-Analysis Procedures

#### Dataset Identification and Selection

Potential datasets were identified in the *Gemma* database using search terms related to rodent SD paradigms (n=106, *search_datasets()* function in package *gemma.R* v.0.99.30), and narrowed using standardized inclusion/exclusion criteria (**Figure 1**, criteria: dx.doi.org/10.17504/protocols.io.j8nlk84jxl5r/v1, completed by researcher CAR). Datasets that lacked relevant SD differential expression results in *Gemma* or samples from the murine or rat cerebral cortex were removed. An additional dataset was removed due to brain collection occurring multiple days (192 hrs) following SD. Following exclusions, 8 relevant datasets remained containing 18 SD interventions (SD vs. control contrasts) (**Table 1**). At this stage, all inclusion/exclusion decisions were reviewed by a second researcher (MHH) and final decisions discussed with the full 2022 *Brain Data Alchemy Project* cohort (CAR, MHH, EF, JX, EH, DMN, AS).

**Figure 1.**
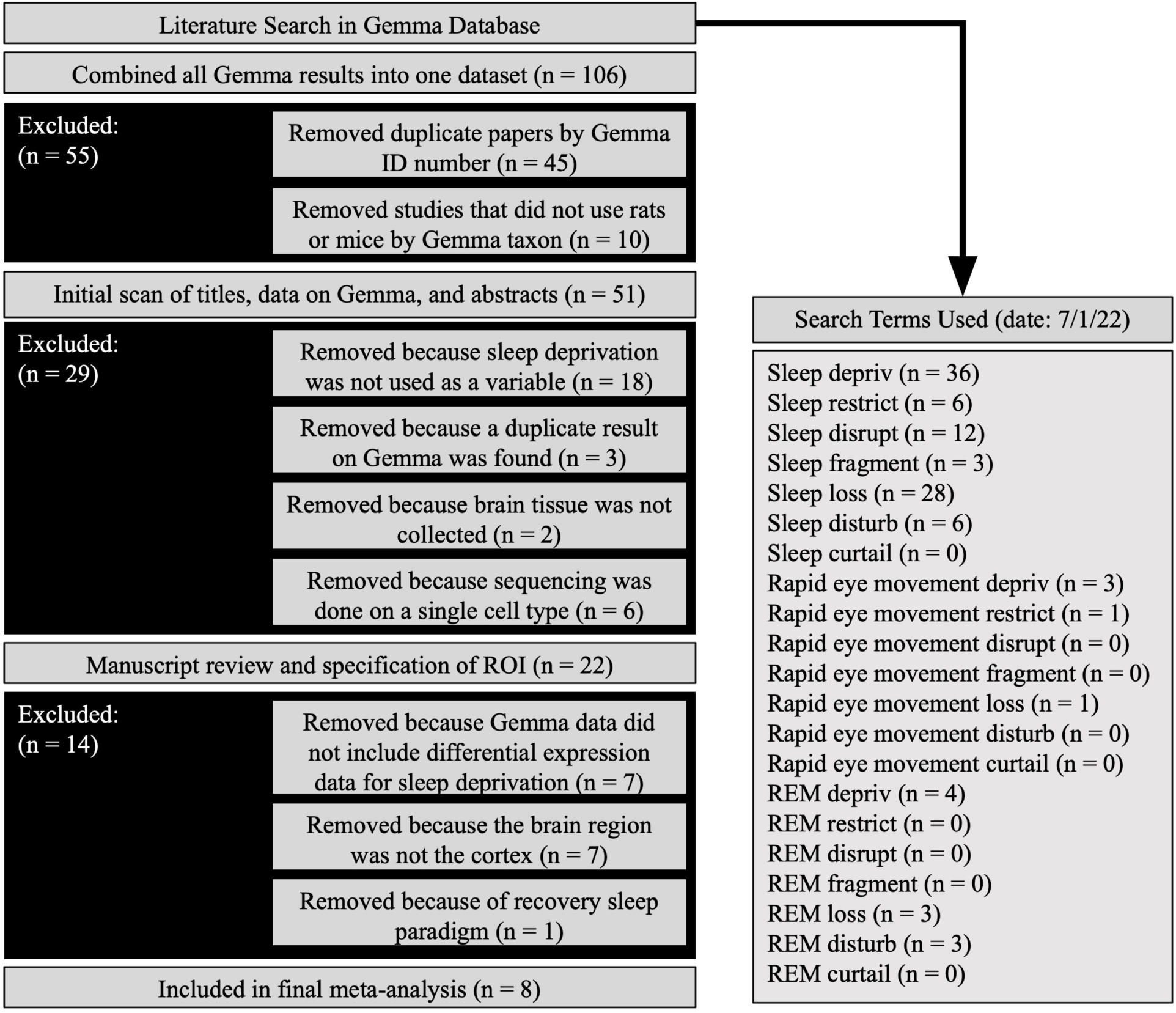
Diagram illustrating the dataset search terms and selection criteria used for the meta-analysis. Abbreviations: ROI (Region of Interest), n (number of datasets)

**Table 1.** Overview of the datasets included in the meta-analysis and in the validation analysis. The table lists the Gene Expression Omnibus (GEO) ID for the dataset, sample size (n) for the subset of the dataset used in the differential expression analysis, the species/strain, sex, and age for the subjects, the authors and year of publication (Bjorness et al., 2020; Diessler et al., 2018; Gerstner et al., 2016; Guo et al., 2019; Hinard et al., 2012; Ingiosi et al., 2019; Mackiewicz et al., 2007; Muheim et al., 2019; Orozco-Solis et al., 2017), the brain region sampled, and transcriptional profiling platform. The table also lists the scope of the transcriptional profiling data included in the current analysis, including the number of probesets (microarray), the average number of aligned RNA-Seq reads and the range of aligned reads across samples (along with the average % of reads uniquely aligned and %GC content), and the number of genes that were represented in the final analysis following quality control procedures. Study details include the SD paradigm and specific experimental groups included in the statistical contrasts extracted for inclusion in the meta-analysis. <u>Abbreviations</u>: wk (week), mo (month), bp (base pairs), M (million), GC (proportion of guanine and cytosine bases), h (hour), SD (sleep deprivation), RS (recovery sleep), Ctrl (control)

| Dataset ID | n | Species | Sex | Age | Authors | Year | Region | Platform | Scope of Profiling | Type of Sleep Dep | Comparisons of Interest ("Contrasts") |
| --- | --- | --- | --- | --- | --- | --- | --- | --- | --- | --- | --- |
| GSE6514 | 45 | Mouse (C57BL/6J) | All male | Adult: 10 wk ± 1 wk | Mackiewicz et al. | 2007 | Cerebral Cortex | Affymetrix GeneChip Mouse Genome 430 2.0 Array | Probesets: 45,101, Final genes: 18,536 | Gentle Handling | 6h SD vs. Ctrl<br>9h SD vs. Ctrl<br>12h SD vs. Ctrl |
| GSE33491 | 9 | Mouse (C57BL/6J) | All male | Not reported | Hinard et al. | 2012 | Cerebral Cortex | Affymetrix Mouse Exon 1.0 ST Array | Probesets: 22,357, Final genes: 18,785 | Unknown | 6h SD vs. Ctrl<br>6h SE + 18h RS vs. Ctrl |
| GSE78215 | 65 | Mouse (C57BL/6J) | All male | Adult: 2 mo | Gerstner et al. | 2016 | Cerebral Cortex | Affymetrix Mouse Gene 2.1 ST Array | Probesets: 41,345, Final genes: 24,027 | Gentle handling | 6h SD + 1h RS vs. Ctrl<br>6h SD + 2h RS vs. Ctrl<br>5h SD + 3h RS vs. Ctrl<br>5h SD + 6h RS vs. Ctrl |
| GSE93041 | 9 | Mouse (C57BL/6J) | All male | Adult: 6 mo | Orozco-Solis et al. | 2017 | Anterior Cingulate Cortex | Affymetrix Mouse Gene 2.0 ST Array | Probesets: 41,801, Final genes: 24,036 | Constant movement | 12h SD vs. Ctrl |
| GSE113754 | 20 | Mouse (C57BL/6J) | All male | Adult: 9 wk | Ingiosi et al. | 2019 | Prefrontal Cortex | 100 bp paired-end RNA-seq, Illumina HiSeq 2500 (HiSeq SBS kit v4) | Aligned reads: 42M (range: 28-77M, 86% uniquely aligned, 48% GC), Final genes: 22,492 | Gentle handling | 5h SD vs. Ctrl |
| GSE128770 | 108 | Mouse (C57BL/6J) | All male | Adult: 2-4 mo; Older Adult: 18-20 mo | Guo et al. | 2019 | Medial Prefrontal Cortex | 100 bp single-end RNA-seq, Illumina HiSeq 4000 (Illumina TruSeq Stranded mRNA kit) | Aligned reads: 21M (range: 16-28M, 86% uniquely aligned, 43% GC), Final genes: 20,017 | Gentle handling | 3h SD vs. Ctrl<br>6h SD vs. Ctrl<br>9h SD vs. Ctrl<br>12h SD vs. Ctrl |
| GSE132076 | 4 | Mouse (C57BL/6J) | All male | Adult: 20 wk ± 2.7 wk | Muheim et al. | 2019 | Cerebral Cortex | 100 bp single-end RNA-seq, Illumina HiSeq 2500 (kit not specified) | Aligned reads: 37M (range: 31-48M, 80% uniquely aligned, 46% GC), Final genes: 24,460 | Gentle handling | 4h SD vs. Ctrl |
| GSE144957 | 33 | Mouse (C57BL/6J) | All male | Adult: 8-19 wk | Bjorness et al. | 2020 | Frontal Cortex | 150 bp paired-end RNA-seq, Illumina NextSeq 500 (TrueSeq Stranded mRNA Library Prep) | Aligned reads: 53M (range: 19-165M, 78% uniquely aligned, 44% GC), Final genes: 23,358 | Gentle handling | 6h SD vs. Ctrl<br>4h SD + 2h RS vs. Ctrl |
| **Validation Analysis:**<br>GSE114845 | 86 (pools of 2-3) | Mouse (full genetic panel) | All male | Adult: 11-14 wk | Diessler et al. | 2018 | Cerebral Cortex | 100 bp single-end RNA-Seq, Illumina HiSeq 2500 (HiSeq SBS Kit v3) | Aligned reads: 34M (range: 28-44M, 83% uniquely aligned), Final genes: 19,797 | Gentle Handling | 6h SD vs. Ctrl |

##### Overview of Selected Datasets

The eight selected datasets included protocols with SD durations that varied between 3-12 hrs. Three datasets had protocols that included recovery sleep (RS), defined as the period between an SD treatment and sacrifice of the animal during which time the animal was freely allowed to sleep, varying between 1-18 hrs. The final collective sample size was *n*=293 mice, with sufficient power (80%) to detect medium effect sizes using a traditional alpha (0.05).

##### Result Extraction

The differential expression results for each of the relevant 18 statistical contrasts (SD vs. control group comparisons) were imported into R using Gemma’s API (package *gemma.R* v.0.99.30). Rows lacking unambiguous gene symbol annotation were excluded, and gene-level average effect sizes defined as log2 fold change (Log2FC) and their respective standard errors and sampling variances (SV) were calculated and aligned across datasets by gene symbol.

##### Meta-Analysis

To perform the meta-analysis, we fit a random effects model to the SD Log2FC values and accompanying SV for each gene that was represented in at least 13 of the 18 contrasts (*n*=16,290 genes). Random effects modeling was chosen to account for the heterogeneous methods and sample characteristics across the included studies that might introduce variability among their true effects. To fit the random effects model, we used the function *rma()* from the *metafor* package (v.3.4.0, (Viechtbauer, 2010)), which treats this heterogeneity as purely random (normally distributed), estimated using restricted maximum-likelihood estimation (REML). The function uses the inverse-variance method, which treats the precision of each study’s estimated effect as inversely related to the study’s sampling variance. Due to limited dataset availability, we used a simple, intercept-only model as our main outcome. As a secondary (exploratory) outcome, we re-ran the meta-analysis with SD duration included as a numeric predictor (centered on average duration) and recovery sleep (RS) included as a factor (intercept=no RS). The resulting p-values were corrected for false discovery rate (FDR) using the Benjamini-Hochberg method (*multtest* package: v.2.8.0, (Pollard et al., 2005)).

##### Functional Ontology

To identify functional patterns, *fast Gene Set Enrichment Analysis* (*fGSEA*) was performed on the meta-analysis results (Sergushichev, 2016), ranked by either estimated Log2FC (directional analysis) or the absolute value of the estimated Log2FC (non-directional analysis), using the *Brain.GMT* gene set database (v1: (Hagenauer et al., 2024b)) and associated example code (https://github.com/hagenaue/Brain_GMT). Brain.GMT is a curated database of gene sets related to nervous system function, tissue, and cell types, packaged with traditional gene ontology gene sets.

#### Result Validation: Comparison with GSE114845

##### Overview of GSE114845

Our meta-analysis focused exclusively on public datasets represented in the *Gemma* database and re-analysis pipeline. This focus precluded the use of a large public RNA-Seq dataset (*GSE114845*, (Diessler et al., 2018)) that was not available on Gemma. GSE114845 examined the effects of SD in mice using a genetic reference population. This dataset included 86 publicly-available RNA-Seq samples (*n*=43 SD, *n*=43 CTRL) from the cerebral cortex, with each sample typically representing pooled RNA from 2-3 biological replicates (*i.e.,* mice from the same line/strain and experimental condition, full *n*=222 mice). As this sample size was on par with our full meta-analysis (*n*=293), we decided to use GSE114845 as independent validation for our meta-analysis results.

##### Re-analysis of GSE114845

It was necessary to re-analyze GSE114845, as the full differential expression results were not released at publication (Diessler et al., 2018). Aligned read counts were generated using the standardized *Archs4* pipeline (Lachmann et al., 2018), which uses *Kallisto* (Bray et al., 2016) to align public mouse RNA-Seq samples against an updated genome (GRCm39) with Ensembl annotation (Ensembl 107). Within R (v.4.5.0), these counts were downloaded using the *rhdf5* package (v.2.53.1, (Fisher et al., 2025)) and data relevant to the cortical samples extracted using *h5read()* function. Additional sample metadata (condition, genotype, tissue) was imported using the *getGEO()* function in the *GEOQuery* package (v.2.77.0, (Davis and Meltzer, 2007)).

We then followed a quality control and analysis procedure as similar as possible to that used by the original publication (Diessler et al., 2018). We filtered out rows (genes) with a mean raw count <10. Library size varied widely by genotype, so TMM normalization was applied using the *calcNormFactors()* function (package: *edgeR,* v.4.7.2, (Chen et al., 2025)) to provide scale factors for estimated relative RNA production levels at the point of calculating counts per million (cpm) or log2cpm (function *cpm()* in *edgeR* (Robinson and Oshlack, 2010)). Differential expression was calculated using a simple model containing only SD treatment – we did not include genotype as a co-variate to avoid model overfitting. Differential expression was calculated using the *limma-voom* pipeline (*limma* package, v.3.65.1, (Law et al., 2014; Ritchie et al., 2015)), with an empirical Bayes correction (function *eBayes()* package *limma*) and FDR correction using the Benjamini-Hochberg method.

##### GSE114845 vs. Meta-Analysis Comparison

For all genes present in both GSE114845 and the meta-analysis output, we determined consistency in both the direction of effect of SD and survival of FDR correction (FDR<0.05). The effect of SD (Log2FC) measured in our meta-analysis and GSE114845 was compared using simple linear regression (*stats* base package: *lm()*) and Spearman’s rank correlation (*stats* package: *cor.test()*) using both the top DEGs from our meta-analysis (FDR<0.05) and all genes present in both datasets. Note that it was important to consider the congruence of the direction of effect between the two datasets and not just validate based on statistical significance as the majority of the cortical transcriptome was found to be differentially expressed in response to SD in *GSE114845* (63% FDR<0.05 in our hands, 78% in the original publication (Diessler et al., 2018)).

## Results

### Planned Meta-Analysis: Overview

Eight datasets examining the effects of SD on the cortex survived our inclusion/exclusion criteria (Bjorness et al., 2020; Gerstner et al., 2016; Guo et al., 2019; Hinard et al., 2012; Ingiosi et al., 2019; Mackiewicz et al., 2007; Muheim et al., 2019; Orozco-Solis et al., 2017), and the differential expression results from eighteen relevant SD vs. control statistical contrasts from those datasets were extracted for inclusion in our meta-analysis (collective *n*=293). Of the 16,290 genes included in the meta-analysis, 16,255 produced stable meta-analysis estimates (full results: **Table S1)**. Our meta-analysis revealed 182 differentially expressed genes (“DEGs”: FDR<0.05), 104 of which were upregulated and 78 downregulated. To illustrate these effects, we have provided example forest plots for two of the top DEGs (glucocorticoid receptor *Nuclear Receptor Subfamily 3 Group C Member 1 (Nr3c1)* and immunoglobulin *CD7 Molecule (Cd7)*: **Figure 2**, other examples: **Figure S1&S2**). We have also provided a hierarchically-clustered heatmap illustrating effects (Log2FC) for the top 50 DEGs (**Figure 3A)**, and a volcano plot summarizing the meta-analysis results overall (**Figure 3B).**

**Figure 2.**
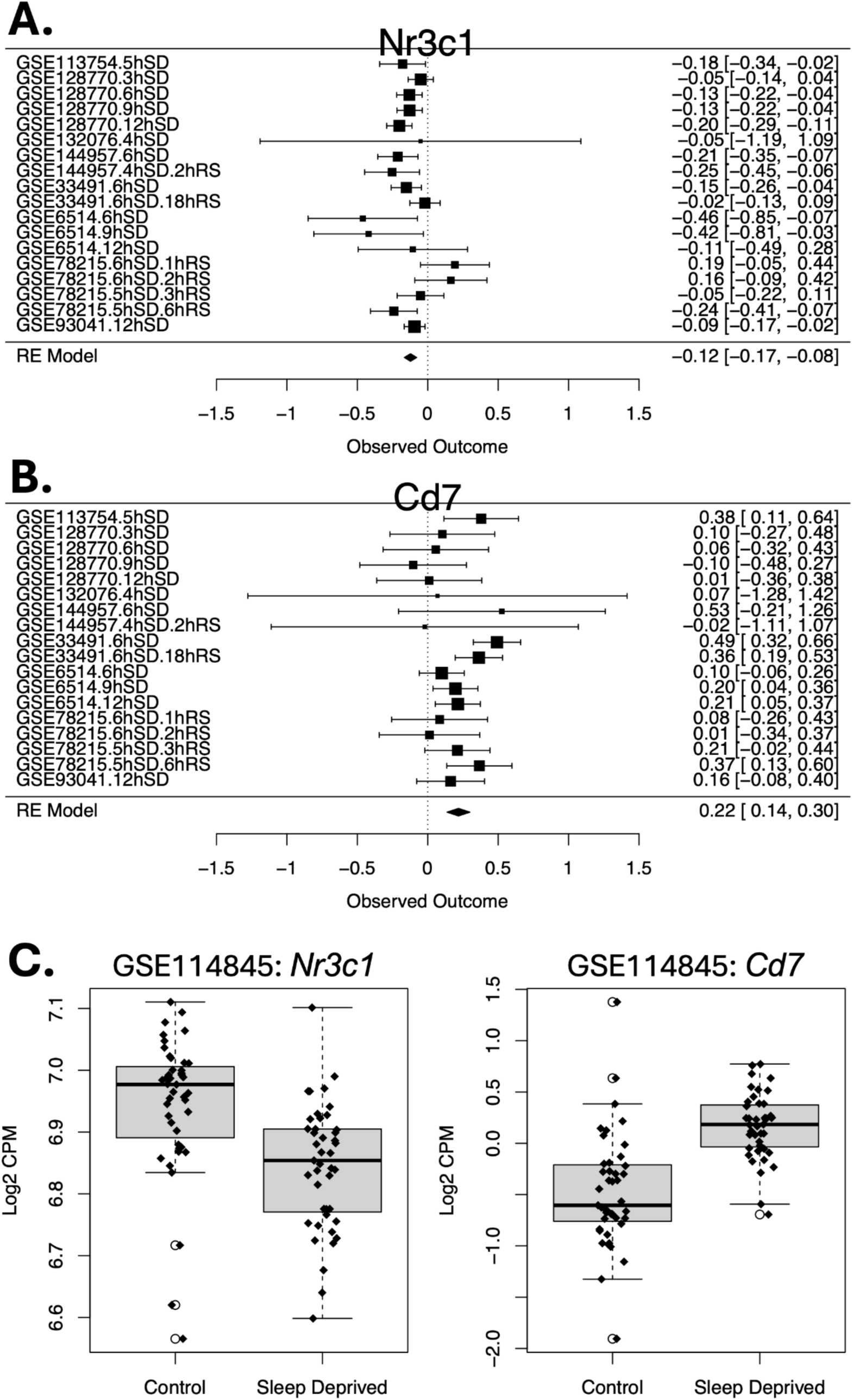
Example forest plots that show genes that were consistently differentially expressed in the murine cortex across SD paradigms and experiments in our meta-analysis of public datasets (collective n=293) and validated using an independent large RNA-Seq dataset (GSE114845). Rows illustrate the effect sizes (squares: SD vs. control Log2FC) for each of the SD vs. control contrasts in the individual datasets and for the random effects meta-analysis model (“RE Model”), accompanied by 95% confidence intervals (whiskers). Each study is named using the Gene Expression Omnibus accession number (GSE#), the duration of SD in hours (h), and, if relevant, the duration of recovery sleep (RS) in hours (hr). Note that the confidence intervals for the individual study contrasts are partially dependent on sample size – this is why GSE132076 (n=4) has very large confidence intervals in all forest plots. **A.** Nuclear Receptor Subfamily 3 Group C Member 1 (Nr3c1) was consistently down-regulated across SD paradigms and experiments. Nr3c1 is the glucocorticoid receptor, involved in stress response pathways. **B.** CD7 Molecule (Cd7) was consistently upregulated across SD paradigms and experiments. Cd7 is a member of the immunoglobulin superfamily and plays an important role in immune function. **C.** <u>Validation</u>: Nr3c1 is also down-regulated following 6 hrs of SD in GSE114845 (FDR<0.05). A box plot with overlaid jittered data points illustrates the relationship between Nr3c1 expression (log(2) counts per million (cpm)) and treatment group (control vs. SD). Each of the 86 RNA-Seq samples represents 2-3 biological replicates pooled from the full sample of n=222 mice. boxes = first quartile, median, and third quartile; whiskers = range or 1.5× the interquartile range; open dot = outlier datapoint falling beyond the whiskers of the boxplot. **D.** <u>Validation</u>: Cd7 is also up-regulated following 6 hrs of SD in GSE114845 (FDR<0.05). The box plot follows the plotting conventions of panel C.

**Figure 3.**
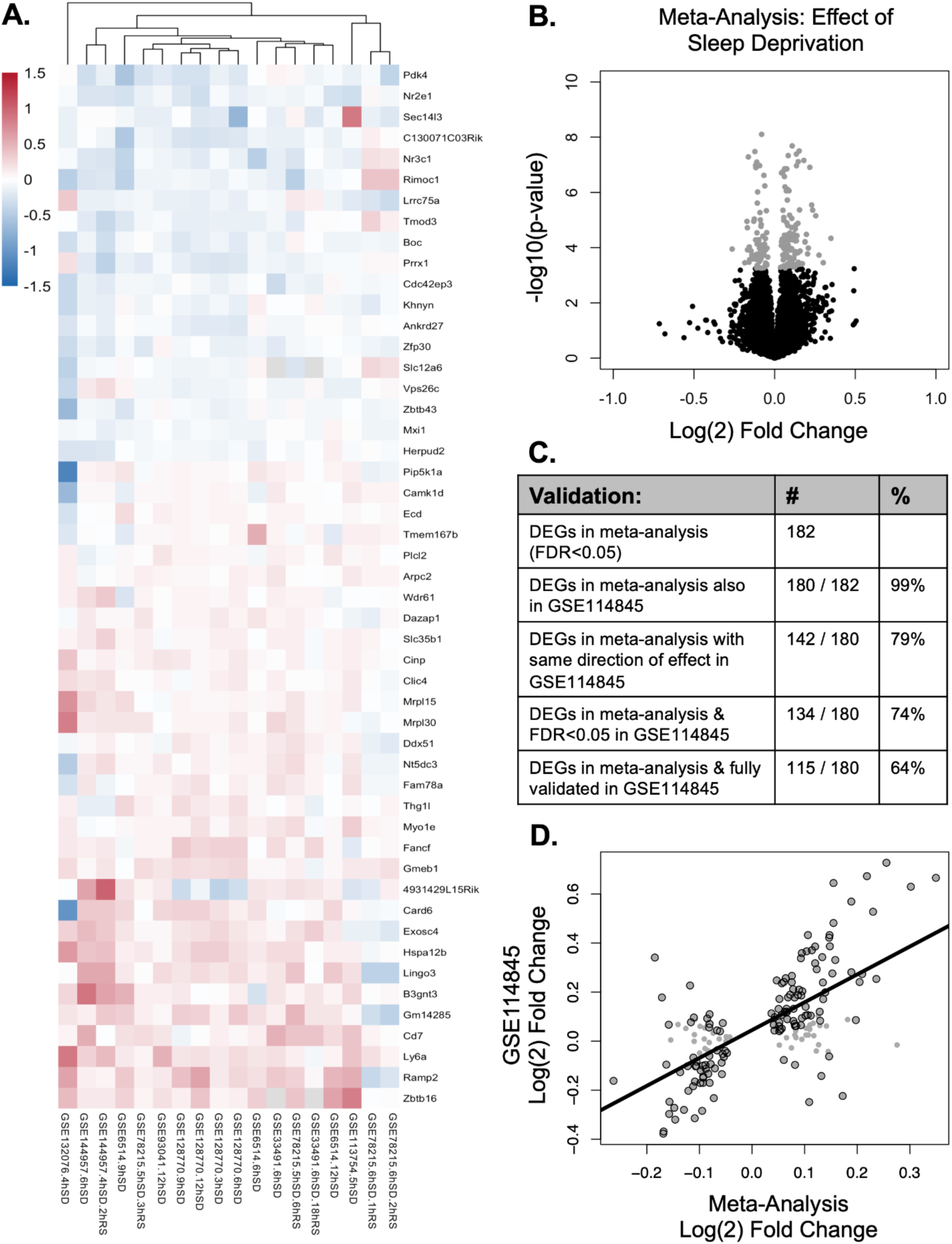
A meta-analysis of public datasets examining the effects of SD on the murine cortex (collective n=293) revealed 182 genes that were consistently differentially expressed across SD paradigms and experiments (“DEGs”: FDR<0.05), most of which showed similar effects in an independent large RNA-Seq dataset (GSE114845). **A.** A hierarchically-clustered heatmap illustrates the effect of SD on gene expression (Log2FC) across each of the datasets and conditions for the top 50 DEGs identified in the meta-analysis. Upregulation following SD is illustrated with red, down-regulation following SD with blue. Genes are identified by official gene symbol. Datasets are identified by Gene Expression Omnibus (GEO) accession number, the duration of SD, and duration of recovery sleep (RS, if included). **B.** A volcano plot illustrating the relationship between the magnitude of the effect of SD (Log2FC) in the meta-analysis and nominal p-value (-log10 transformed, so that the smallest p-values are near the top of the y-axis). The 182 DEGs that survived FDR correction (FDR<0.05) are colored grey. **C.** <u>Result</u> <u>validation</u>: A table illustrating the use of a separate large independent RNA-Seq dataset (GSE114845) as validation for the meta-analysis results. Of the 182 DEGs from the meta-analysis (FDR<0.05), 180 were represented in GSE114845, 115 of which (64%) showed both the same direction of effect in GSE114845 and survived FDR correction (FDR<0.05,“fully validated”). **D.** <u>Result validation</u>: A scatterplot illustrating the positive correlation (Spearman’s rho: 0.611, p<2.00E-16) between the effect of SD on gene expression (Log2FC) estimated within our meta-analysis and the effect of SD on gene expression (Log2FC) from GSE114845. Each grey dot represents a DEG that survived false discovery rate correction in our meta-analysis (FDR<0.05), with a black outline indicating that the DEG also survived FDR correction in GSE114845 (FDR<0.05).

### Result Validation: Comparison with GSE114845

Our meta-analysis focused exclusively on public datasets represented in the *Gemma* database and re-analysis pipeline, precluding the inclusion of a large public SD RNA-Seq dataset (GSE114845, (Diessler et al., 2018): *n*=86 cortical RNA-Seq samples pooled from *n*=222 mice). As an independent validation, we compared our meta-analysis results to the differential expression results from GSE114845 (full re-analysis results: **Table S2)**. When considering the top DEGs from our meta-analysis (n=182, FDR<0.05), we found that the estimated Log2FCs from the meta-analysis strongly correlated with the Log2FCs from GSE114845 (β+/-SE: 1.137+/-0.097, *T*(178)=11.727, *p*<2.00E-16, Spearman’s rho: 0.611, **Figure 3D**). Most of the DEGs from our meta-analysis showed the same direction of effect in GSE114845 (79% congruence: 142 of 180 DEGs represented in GSE114845). Moreover, most of the DEGs that showed the same direction of effect in our meta-analysis and GSE114845 also survived FDR correction in GSE114845 (FDR<0.05) and were deemed fully validated (115/182 DEGs, 64%; **Figure 3C**). For these fully validated DEGs, we have provided the meta-analysis results and GSE114845 differential expression results in **Table 2** and **Table 3**, with the non-validated DEGs found in **Tables S3** and **S4**.

**Table 2.**
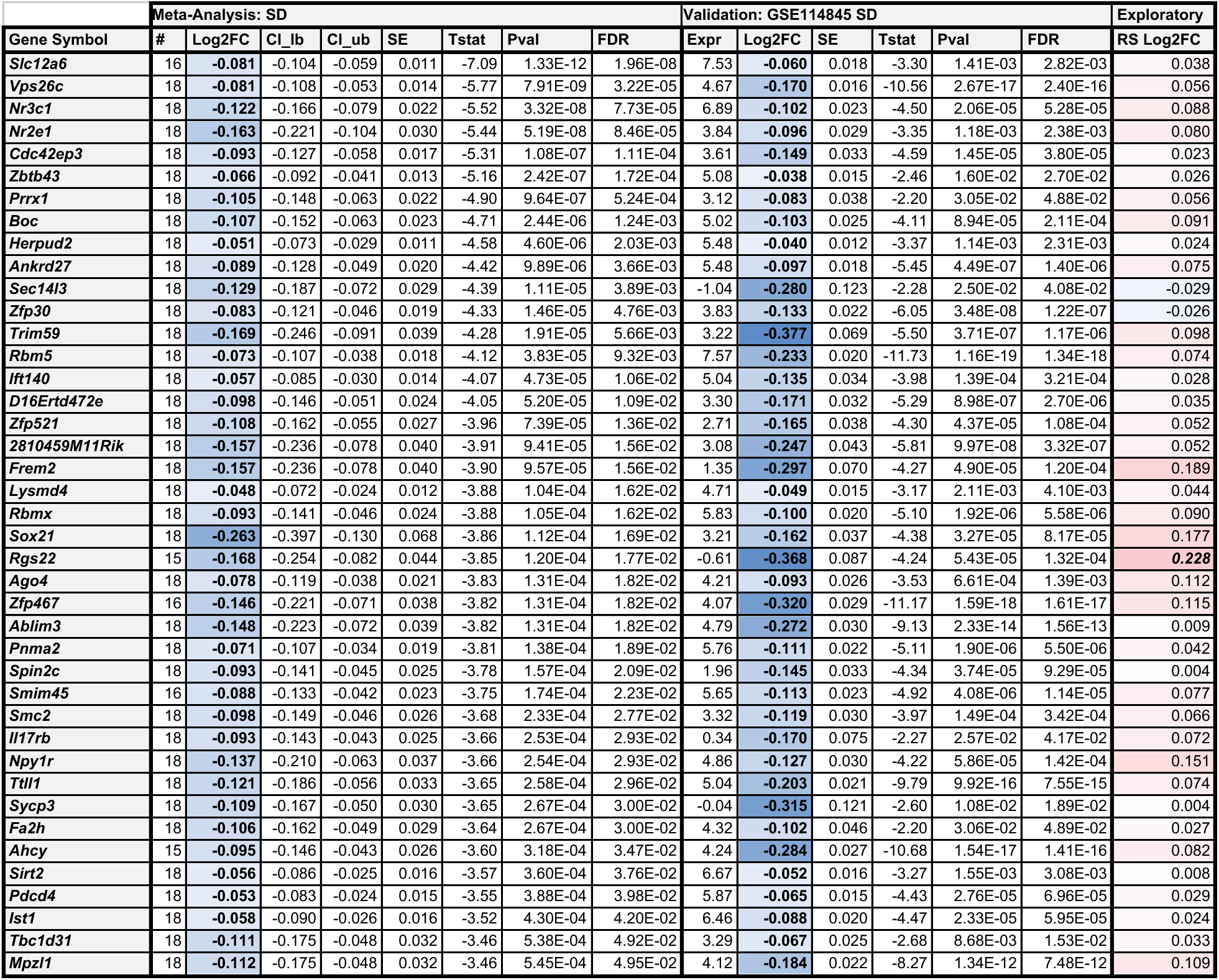
Genes that are downregulated in the murine cortex following SD. The downregulation reported in this table was observed both in our meta-analysis of public transcriptional profiling data (FDR<0.05, collective n=293), and validated by a separate large RNA-Seq study (GSE114845). Genes that showed downregulation in our meta-analysis that were not validated using GSE114845 can be found in **Table S3**. Within the Log(2)Fold Change (Log2FC) columns, blue is used to highlight down-regulation, pink is used to highlight up-regulation, bold text indicates FDR<0.05, and bold/italic text indicates nominal significance (p<0.05). <u>Column</u> <u>definitions</u>: #= Number of differential expression results (SD vs. control statistical contrasts) that contributed to the meta-analysis estimate for that gene; Log2FC=estimated SD vs. control Log(2) Fold Change; CI_lb & CI_ub=Lower and upper bound for the 95% confidence interval for the Log2FC; SE=Standard error for the Log2FC, Tstat=T-statistic, Pval=Nominal p-value, FDR=False discovery rate (q-value or adjusted p-value), Expr=Average Log2 expression (log2 counts per million: lcpm) for the gene, RS Log2FC: estimated moderating effect (Log(2) Fold Change) of recovery sleep (RS) on the effect of SD within an exploratory meta-analysis – note that the effect of RS is almost always in the opposite direction of the effect of SD.

**Table 3.**
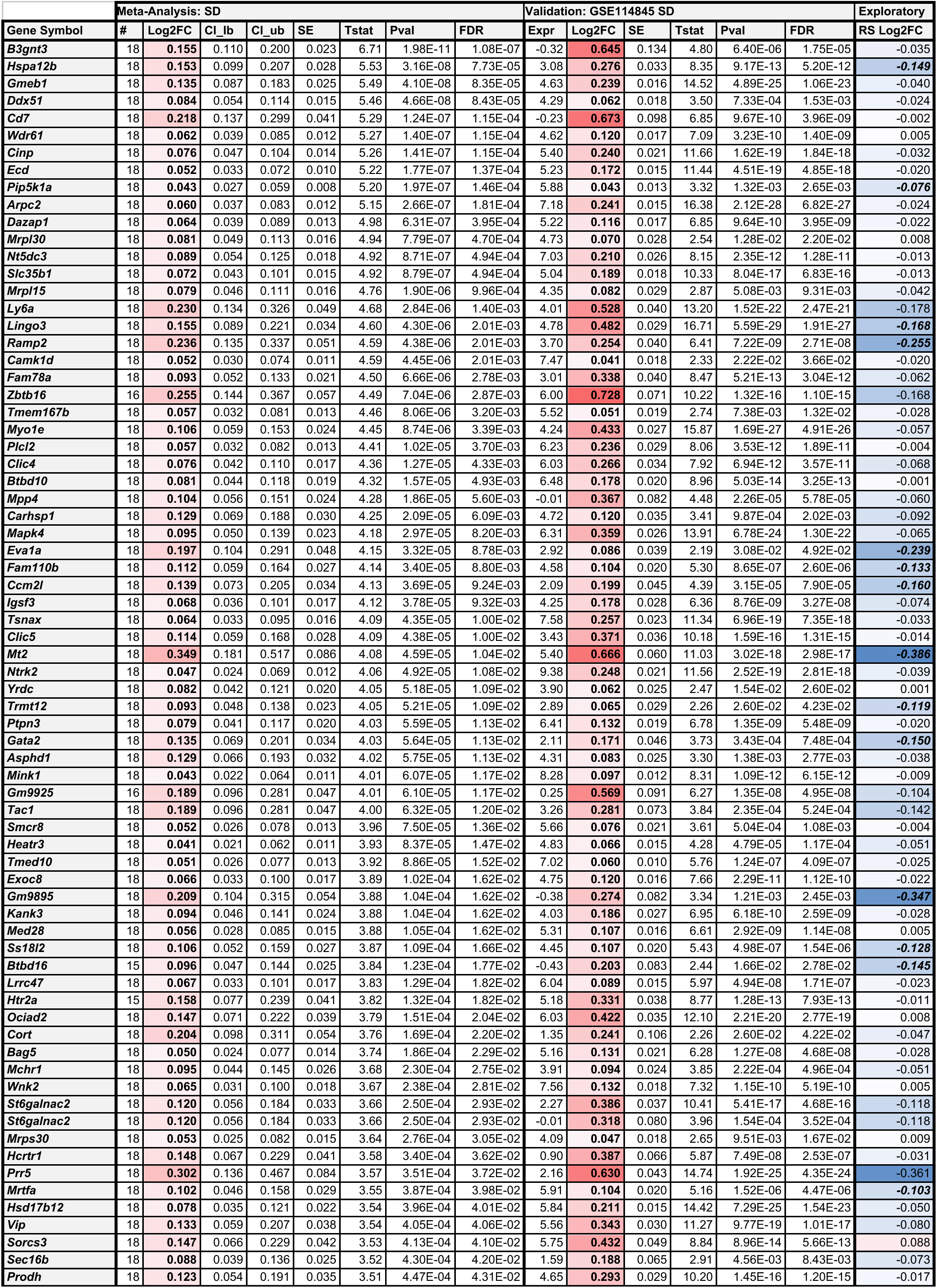
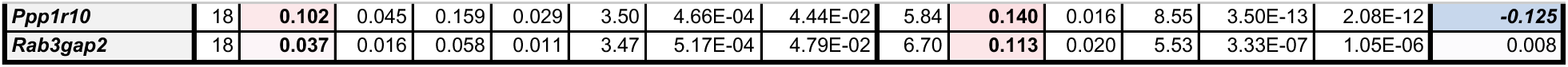
Genes that are upregulated in the murine cortex following SD. The upregulation reported in this table was observed both in our meta-analysis of public transcriptional profiling data (FDR<0.05, collective n=293), and validated by a separate large RNA-Seq study (GSE114845). Genes that showed upregulation in our meta-analysis study that were not validated using GSE114845 can be found in **Table S4**. Column definitions are the same as Table 2.

#### Functional Ontology

fGSEA revealed biological pathways enriched with differential expression following SD, as identified by our meta-analysis. Of the 10,436 gene sets included in the fGSEA output, 236 gene sets were enriched with differential expression (FDR<0.05, examples: **Table 4**, full results: **Table S5**), most of which were downregulated (200 of 236). Most of the down-regulated gene sets (155 gene sets) were dominated by a correlated cluster of leading edge genes that included the DEGs *Nuclear Receptor Subfamily 3 Group C Member 1* (*Nr3c1)*, which is the glucocorticoid receptor, and *Nuclear Receptor Subfamily 2 Group E Member 1* (*Nr2e1)*, as well as many immediate early genes (e.g., *Fos Proto-Oncogene, AP-1 Transcription Factor Subunit* (*Fos)*) which were not individually significantly differentially expressed in the meta-analysis. These down-regulated gene sets were predictably involved in the stress response and immediate early gene responses, but also pathways associated with transcription regulation, plasticity, growth and development, and vasculature. When considering only down-regulated gene sets driven by DEGs that were validated in GSE114845 and not dominated by immediate early genes (23 gene sets), the predominant theme was growth and development (17 gene sets), including pathways related to progenitor cells, epithelial development, neuron differentiation, and embryonic morphogenesis. These gene sets often included DEGs *Nr2e1, FRAS1 Related Extracellular Matrix 2 (Frem2), Intraflagellar Transport 140 (Ift140), and Paired Related Homeobox 1 (Prrx1)*.

**Table 4.**
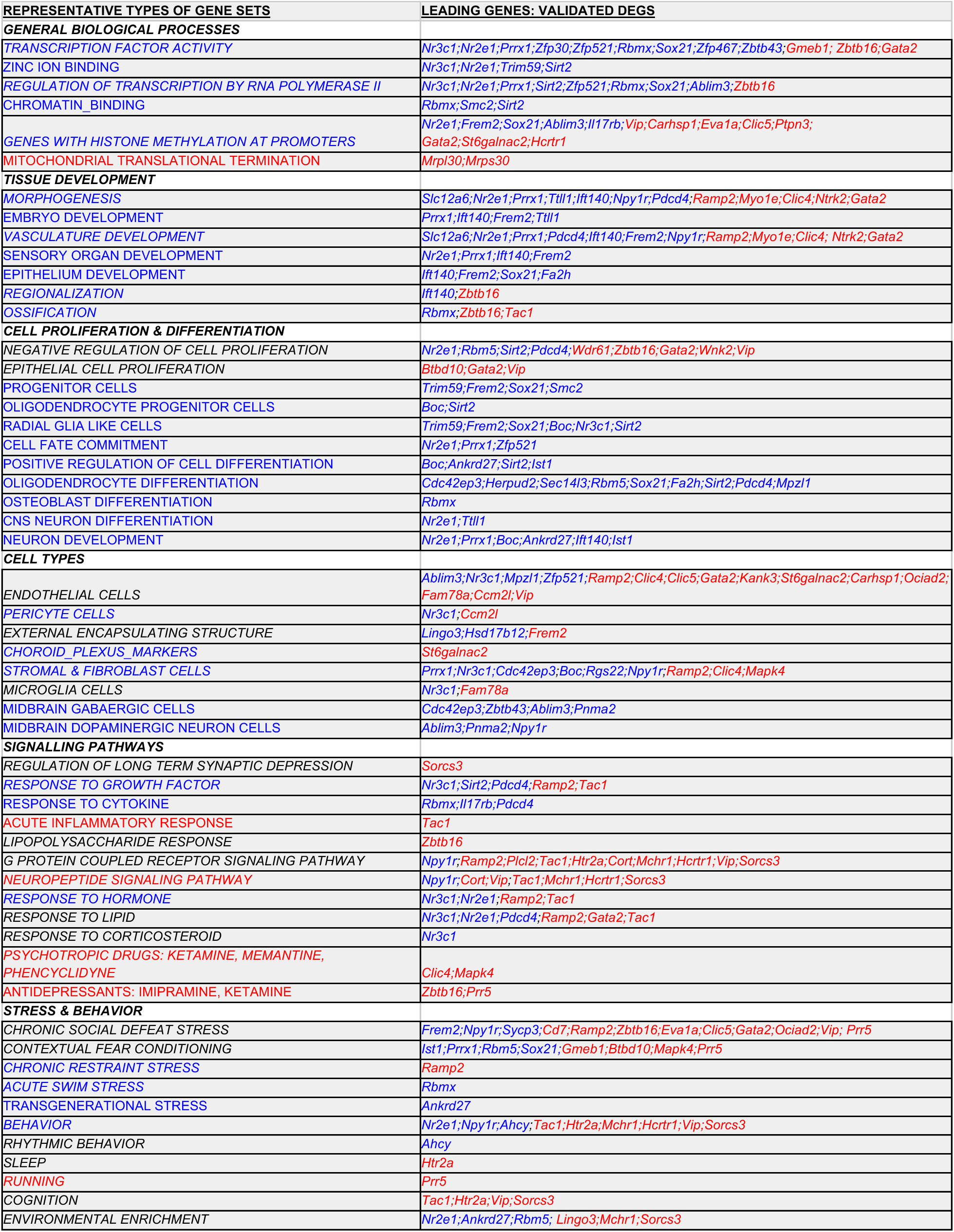
SD affects cortical gene expression within pathways associated with transcription regulation, development, cell proliferation and differentiation, cell signalling, stress, and behavior. Included in the table are representative types of gene sets that were enriched with differential expression (FDR<0.05) and associated with validated DEGs. Blue text indicates enrichment of down-regulation following SD within a type of gene set, red text indicates enrichment with up-regulation following SD, black text indicates that there was a contradictory direction of effect amongst related gene sets or that the gene set was only significant in the non-directional analysis, and italic font indicates enrichment with differential expression in the non-directional analysis. The validated DEGs associated with each type of gene set are listed using official gene symbol, with font colors indicating their direction of effect (blue=down-regulated with SD, red=up-regulated with SD).

In contrast, upregulated gene sets were associated with pathways related to adaptation challenges, such as stress and inflammation, with leading edge genes including known mediator DEG *CD7 Molecule* (*Cd7:* in 15 gene sets), *Zinc Finger And BTB Domain Containing 16* (*Zbtb16)* and *Proline Rich 5* (*Prr5)*. In parallel, there was upregulation of mitochondrial translation (including DEGs *Mitochondrial Ribosomal Protein L30* (*Mrpl30)* and *Mitochondrial Ribosomal Protein S30* (*Mrps30)*). Notably, pathways related to neuropeptide signalling were also upregulated, including the DEGs *Cortistatin* (*Cort*)*, Vasoactive intestinal polypeptide (Vip),* and *Tachykinin Precursor 1 (Tac1),* and receptors *Melanin-concentrating hormone receptor 1 (Mchr1)* and *Hypocretin receptor 1 (Hcrtr1),* as well as gene sets related to psychotropic drugs and antidepressants.When considering only upregulated gene sets driven by DEGs that were validated in GSE114845 (19 gene sets), stress, inflammation, and mitochondrial translation continued to be themes, as well as neuropeptide signalling.

To account for gene sets that may include both up– and down-regulated genes, we ran a nondirectional fGSEA that confirmed and expanded on our initial analysis. Of the significant nondirectional gene sets (158 gene sets with FDR<0.05), pathways were largely reflective of those seen in our directional fGSEA (*e.g.,* transcription regulation, growth and development, vasculature, plasticity, stress, neuropeptide signalling, and microglia). Importantly, we saw additional enrichment in pathways related to sleep and rhythms, driven by DEGs *adenosylhomocysteinase (Ahcy)* and serotonin receptor *5-hydroxytryptamine receptor 2A* (*Htr2a),* and in gene sets broadly associated with behavior and cognition, and a comparatively limited number of pathways related to inflammation.

### Exploratory Meta-Analysis

Within our planned meta-analysis, the estimated effects of SD were generally small in magnitude (*abs*(Log2FC)<0.25, **Figure 3B**), which could imply a dilution due to heterogeneity across studies. When examining differential expression for the top DEGs within each of the individual experiments and conditions (**Figure 3A**) we did not observe any obvious clustering related to potential sources of heterogeneity, such as SD duration, the addition of recovery sleep (RS), transcriptional profiling platform, or cortical region. However, this lack of heterogeneity amongst the top DEGs might be expected, as our meta-analysis was designed to detect consistency across studies.

Therefore, we ran a follow-up exploratory meta-analysis to examine whether the SD differential expression results were modulated by SD duration or the addition of RS following SD, as theory would dictate (full results: **Table S6**). For the 8 datasets (n=18 contrasts) included in the meta-analysis, SD duration ranged from 3-12 hours, and 3 datasets (n=6 contrasts) included RS (ranging from 1-18 hrs). We found that SD duration did not significantly modulate SD effects. The addition of RS, however, induced a significant reversal of SD differential expression in a small set of genes with diverse functions: *Major urinary protein 3* (*Mup3)*, *doublecortin like kinase 1 (Dclk1)*, *aminoadipate aminotransferase* (*Aadat),* and *PRKC, apoptosis, WT1, regulator* (*Pawr)* (example forest plot: **Figure 4A**). Moreover, most of the 182 genes that showed an effect of SD in the planned meta-analysis (FDR<0.05) showed an effect of RS that was in the opposite direction (158 genes or 87%, **Figure 4D**). These reversals were not significant (FDR<0.05), but 21 (11%) showed nominal significance (p<0.05, see **Tables 2&3, Tables S3&S4**). A similar pattern was not observed for SD duration (**Figure S3**).

**Figure 4.**
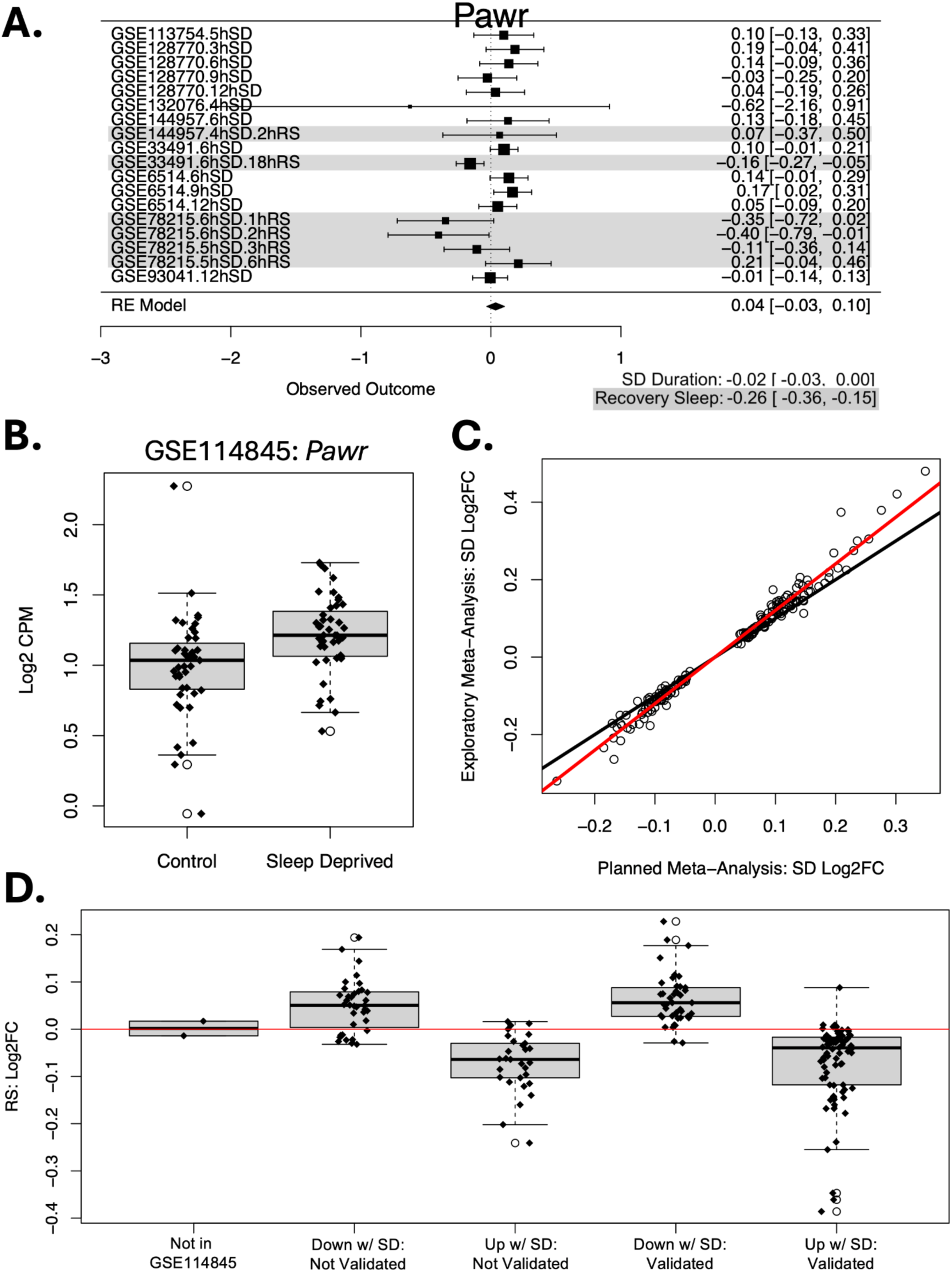
<u>Exploratory:</u> Recovery sleep (RS) may reverse the effect of SD on gene expression. **A.** Example forest plot: Pro-Apoptotic WT1 Regulator (Pawr) was upregulated with SD (RE Model: “Intercept”) in a manner that was not significantly modulated by SD Duration (RE Model: “SD Duration”) but appeared reversed by RS (highlighted gray, RE Model: “Recovery Sleep”). Pawr promotes apoptosis, which is programmed cell death. The forest plot follows the plotting conventions of **Figure 2**. **B.** <u>Validation</u>: Pawr was similarly up-regulated following 6 hrs of sleep deprivation in GSE114845 (FDR<0.05). A box plot with overlaid jittered data points illustrates the relationship between Pawr expression (log(2) counts per million (cpm)) and treatment group (control vs. SD). The box plot follows the plotting conventions of **Figure 2**. **C.** The effect sizes (Log2FC) estimated within our exploratory meta-analysis that controlled for RS and SD duration (y-axis) tended to be slightly larger in magnitude than the effect sizes (Log2FC) estimated in our planned meta-analysis that did not control for RS (x–axis). This pattern suggests that the inclusion of studies with RS diluted effect sizes in our planned meta-analysis. The scatter plot illustrates this linear relationship, focusing on the 182 genes that showed an effect of SD in the planned meta-analysis (FDR<0.05), with the red line showing the slope (β=1.20+/-0.013 SE) and the black line indicating the hypothetical slope if the Log2FC estimates from the two meta-analyses were equivalent (1:1). **D.** Most of the 182 genes that showed an effect of SD in the planned meta-analysis (FDR<0.05) – including both the DEGs that were fully validated using GSE114845 and those that were not – showed an effect of RS (Log2FC: y-axis) that was in the opposite direction of the effect of SD in the planned meta-analysis. None of these effects were significant (FDR<0.05), but 21 (11%) showed nominal significance (p<0.05, see **Tables 2&3, Tables S5&S6**).

Controlling for these sources of heterogeneity appeared to improve our ability to detect DEGs for SD, with 326 genes now showing significant SD effects (FDR<0.05). When focusing on the 182 genes that showed an effect of SD in the planned meta-analysis (FDR<0.05), we noticed that the SD effect sizes (Log2FC) estimated within our exploratory meta-analysis tended to be slightly larger (**Figure 4C**, slope of linear relationship: β=1.20+/-0.013 SE, R^2^=0.98, T(180)=92.5, p<2E-16), as would be expected if including studies with RS in our planned meta-analysis had diluted SD effect size estimates.

## Discussion

Investigating the effects of sleep deprivation (SD) on the brain is important due to the prevalence of sleep disorders and sleep disruption (Li et al., 2018; Patel et al., 2010). Our meta-analysis revealed that SD has varied and multi-directional effects on gene expression in the mouse cerebral cortex that are consistent and reproducible across laboratories and conditions, and that appear to be reversible following recovery sleep (RS). We found an enrichment of differential expression within many functional gene sets, with downregulation related to the stress response, vasculature, plasticity, growth and development, and upregulation related to stress, inflammation, and neuropeptide signalling. These results confirm and expand upon many themes identified in earlier transcriptomic reviews and meta-analyses (*e.g.,* (Cirelli and Tononi, 2001; Elliott et al., 2014; Giannos et al., 2022; Wang et al., 2010)). We discuss the potential role for these effects in the brains’ response to SD in greater depth below.

### Cortical neuropeptide signalling following SD

Reassuringly, our meta-analysis confirmed an upregulation of multiple genes related to neuropeptide signalling in response to SD that have been previously shown to promote SWS (Cirelli and Tononi, 2000; Elliott et al., 2014), including regulators of the brain derived neurotrophic factor (BDNF) pathway. The release of cortical BDNF in response to waketime neural activity contributes to the homeostatic regulation of SWS (Faraguna et al., 2008). BDNF enhances SWA by strengthening excitatory synaptic connections on pyramidal cells (ElGrawani et al., 2024) and *Cortistatin-*releasing interneurons (Bourgin et al., 2007; de Lecea et al., 1996; Martinowich et al., 2011). These effects depend on BDNF receptor *Neurotropic receptor tyrosine kinase 2 (Ntrk2*, also known as tyrosine kinase B), which was upregulated following SD, along with sleep-promoting Cortistatin *(Cort)*. *Tachykinin precursor 1* (*Tac1*) was also upregulated by SD and encodes the precursor for substance P. Cortical microinjections of Substance P increase non-rapid eye movement sleep (NREM) sleep and enhance SWA (Shen et al., 2022).

Other upregulated neuropeptide pathways are known to promote rapid eye movement (REM) sleep. *Vasoactive intestinal polypeptide (Vip)* plays an important role in regulation of the sleep/wake cycle (Hu et al., 2011) and REM sleep (Drucker-Colín et al., 1984). *Vip* is expressed in a subset of cortical interneurons, and was upregulated by SD (Elliott et al., 2014), in a manner that may promote compensatory rebounds in NREM sleep and REM sleep (Hu et al., 2011). Likewise, subcortical Melanin-concentrating hormone (MCH) neurons can promote REM sleep (Potter and Burgess, 2022). The upregulation of *Melanin-concentrating hormone receptor 1* (*Mchr1)* in the cortex in response to SD suggests it may mediate homeostatic effects.

Other neuropeptide signalling may be important for maintaining wakefulness during SD. Neurons containing hypocretin 1 and 2 (also known as Orexin A and B) play a critical role in promoting wakefulness and stabilizing sleep and wake states. Disrupted hypocretin function is linked to narcolepsy, and antagonists of hypocretin receptors can treat insomnia (Pizza et al., 2022). *Hypocretin receptor 2 (Hcrtr2,* or OXR2) appears to mediate many of these effects (Pizza et al., 2022), but our meta-analysis showing upregulation of *Hypocretin receptor 1* (*Hcrtr1,* or OXR1) following SD suggests that *Hcrtr1*’s role may be non-negligible. Down-regulation of neuropeptide Y receptor Y1 (*Npy1r*) following SD may also contribute to maintaining wakefulness, as NPY can inhibit hypocretin neurons and promote sleep (Shen et al., 2022). In addition to neuropeptide signalling, upregulation of monoamine signalling via *Serotonin receptor 5-hydroxytryptamine receptor 2A* (*Htr2a*) during SD may further promote wakefulness, as antagonists of this receptor can reduce insomnia (Vanover and Davis, 2010).

### The relationship between SD, neuroplasticity, growth, and vasculature

Prior literature has implicated sleep as a necessary step in brain growth and plasticity. Sleep promotes learning and memory, and accordingly SD can inhibit it (Lyons et al., 2023). This relationship is theorized to be mediated by synaptic changes precipitated by sleep. Under many conditions, wakefulness promotes synaptic strength and sleep allows for synaptic down-selection (Cirelli and Tononi, 2019; Tononi and Cirelli, 2003). Other studies indicate that synaptic potentiation during sleep may also be important for learning (Kreutzmann et al., 2015), or perhaps sequential synaptic potentiation and synaptic downscaling (Frank and Cantera, 2014).

In our results, neuroplasticity and growth-related gene sets were found to be downregulated by SD. Some genes associated with synaptic potentiation (*e.g.,* BDNF receptor *Ntrk2*) were upregulated by SD, but so were genes mediating long-term synaptic depression (*e.g., Sortilin-Related VPS10 domain-containing receptor 3* (*Sorcs3*) (Breiderhoff et al., 2013)) in a manner that may inhibit BDNF signalling (Subkhangulova et al., 2018). Moreover, we did not observe an induction of immediate early genes following SD, which is thought to mediate synaptic effects (*e.g.,* (ElGrawani et al., 2024; Elliott et al., 2014; Wang et al., 2010), instead observing non-significant down-regulation. As our findings were derived from bulk tissue samples from a variety of cortical regions and experimental paradigms, these contradictions could be due to SD causing effects on synaptic plasticity that vary by synapse type and location (Weiss and Donlea, 2021) or circadian timing (Frank and Cantera, 2014).

Beyond synaptic plasticity, we also saw down-regulation within a variety of gene sets associated with cell proliferation, morphogenesis, growth, and development. These findings are interesting – although sleep is strongly tied to growth-related processes, including nocturnal secretion of growth hormone (Sassin et al., 1969) and child brain development (Jenni and Carskadon, 2007), effects of SD on brain cell proliferation, cell survival, and morphology are typically only observed following more extreme, chronic SD protocols (Kreutzmann et al., 2015). Our findings suggest that the differential expression underlying these effects may begin under acute SD conditions, before histological and morphological changes are visible.

Down-regulation in growth-related pathways could also contribute to the down-regulation observed in gene sets related to vasculature and cerebral spinal fluid regulation (endothelial cells, stromal cells, pericytes, fibroblasts, epithelial cells). These findings are provocative, as these cell types play a central role in two important functions of sleep: the maintenance of proper neurovascular coupling, allowing blood flow to increase in response to local neural activity (Rab-Bábel et al., 2025; Schei and Rector, 2011), and brain waste clearance (Zielinski et al., 2016). During SD, prolonged neural activity blunts vasodilatory reactivity, causing this system to become less responsive (Rab-Bábel et al., 2025; Schei and Rector, 2011). Moreover, chronic SD accelerates vasculature aging, causing hypertension, and arteriosclerosis (Mahalakshmi et al., 2020; Medic et al., 2017). We may have observed the groundwork for these longer term effects, including differential expression within osteogenic pathways related to inflammatory vascular calcification (Mahalakshmi et al., 2020; Shen et al., 2025).

### The relationship between SD, stress, inflammation, and allostasis

Similar to previous studies (Mongrain et al., 2010), within our SD meta-analysis we observed both down– and upregulation in many stress-related gene sets. Indeed, down-regulation of the gene encoding the stress hormone glucocorticoid receptor (*Nr3c1*) was one of the most consistent findings identified across studies, and may represent an adaptation to increased circulating glucocorticoid levels (Juszczak et al., 2025). In response to these observations, we systematically compared our list of SD DEGs to a published database (Table S2 in (Juszczak et al., 2025)) overviewing evidence from high-powered studies characterizing the effects of acute and chronic glucocorticoid or stress exposure on brain gene expression (Jaszczyk et al., 2023; Juszczak et al., 2025; Juszczak and Stankiewicz, 2018; Stankiewicz et al., 2022). We found that 78 of our DEGs have consistent evidence linking them to either glucocorticoid or stress effects on the brain. Of these, 57 DEGs showed effects of glucocorticoids or stress that were in the same direction as SD, including both up and down-regulation (**Table S7**).

The strong overlap between the effects of SD and stress signalling on gene expression may represent an experimental artifact, as the protocols used to keep animals awake can be stressful (Nollet et al., 2020). That said, glucocorticoid elevation following short durations of acute SD in rodents (*e.g.,* 5 hrs) is often within the range of what is observed during normal spontaneous behavior (Kreutzmann et al., 2015). Moreover, the studies included in our meta-analysis and validation analysis mostly used the milder method of gentle handling (Nollet et al., 2020). However, as stress responses to gentle handling can be protocol-, strain-, and experimenter-specific, we cannot rule out greater stress induction, especially since none of the studies reported stress hormone levels.

That said, many of the overlapping effects of SD and stress signalling on gene expression could also represent the fact that SD itself is inherently a challenge to homeostasis, and thus SD activates – and can subsequently overload – many allostatic processes that serve to promote stability in response to challenge, including the stress, metabolic, and immune systems (McEwen and Karatsoreos, 2022). Short durations of SD not only increase glucocorticoids, but also blood pressure, appetite, insulin, and pro-inflammatory cytokines, and decrease parasympathetic tone (McEwen and Karatsoreos, 2022). Suggesting a broader allostatic response, many of the effects that we observed are likely to serve a feedback or protective role, such as the down-regulation of the glucocorticoid receptor (*Nr3c1*) and pro-inflammatory cytokine receptor Interleukin 17 Receptor B (*Il17rb*). Upregulation of *Zinc finger and BTB domain containing 16 (Zbtb16)* following SD may also feed back to limit glucocorticoid effects and enhance repressive (e.g., anti-inflammatory) effects (Galuh et al., 2025), protecting against metabolic and mitochondrial dysfunction (Karagiannopoulos et al., 2023). Similarly, upregulation of heat shock protein *Heat Shock Protein Family A (Hsp70) Member 12B* (*Hspa12b*) following SD may protect the blood-brain-barrier from injury and inflammation (Chen et al., 2017; Zhao et al., 2018) and increased glucocorticoid-activated *Metallothionein 2* (*Mt2*) may protect against oxidative stress and inflammation (Wang et al., 2023).

However, it is also well-established that stress disrupts sleep, and stress hormones, such as glucocorticoids and corticotropin-releasing hormone, can increase arousal and wakefulness, reduce NREM, and suppress the homeostatic response to SD (Nollet et al., 2020). Therefore, we cannot rule out the possibility that the overlapping effects of SD and stress on gene expression may actually represent disrupted sleep within previous stress experiments. For example, several of the neuropeptide-related genes discussed earlier showed similar effects in response to both SD and stress signalling, including *Vip, Ntrk2*, and *Mchr1.* As these neuropeptide pathways have well-known roles in the promotion and regulation of sleep it is unlikely that their differential expression following SD is simply a side effect of elevated stress levels. Future studies inducing SD without stress-inducing protocols, such as by directly inhibiting sleep-promoting neurons or activating wake-promoting neurons, may differentiate between these possibilities (Nollet et al., 2020).

### Exploratory analyses examining SD Duration and Recovery Sleep (RS)

Longer periods of SD should produce a greater build-up of homeostatic drive to sleep, producing more dramatic effects on the brain and behavior (Tobler and Borbély, 1990). However, within an exploratory analysis, we found that the duration of the SD protocols did not significantly modulate the effect of SD. This could suggest a plateau in measurable SD effects on cortical expression such that the SD paradigms used in our meta-analysis (ranging from 3-12 hrs) had similar biological consequences. That said, our power to detect relationships between SD duration and gene expression was smaller than our power to detect simple SD effects because it depends on the representation of the exploratory variable in the statistical contrasts. As ten of our contrasts had an SD duration of 5-6 hrs, there may simply not have been sufficient power amongst the remaining 8 contrasts to garner reliable information about the effect of SD duration, especially given other sources of correlated heterogeneity.

RS following SD should relieve homeostatic pressure and reverse many of the effects of SD (Borbély et al., 1981; Tobler and Borbély, 1990). In our exploratory analysis, RS reversed many of the effects of SD on gene expression. However, the effects of RS were only significant for a handful of genes, again most likely due to limited power. Within the exploratory analysis, we also found that the effects of SD tended to be larger than in our planned meta-analysis, and the number of significant SD DEGs doubled. Collectively, this suggests that the inclusion of studies with RS in our planned meta-analysis may have diluted our original effect size estimates.

### Limitations and future directions

There are several limitations to our analysis that are worth noting. First, sleep need and sleep architecture vary across the lifespan in a manner that depends on sex, hormonal and reproductive factors, and health status (Bishir et al., 2020; Hajali et al., 2019; Jenni and Carskadon, 2007; Mahalakshmi et al., 2020). SD has a larger impact on women, causing a faster accumulation of sleep pressure and increased vulnerability to SD-associated inflammation, metabolic and cardiovascular issues (Hajali et al., 2019). Moreover, there is evidence that SD effects on the cortical transcriptome may depend on sex (Shi et al., 2023) and age (Guo et al., 2019). Therefore, it is unfortunate that every study that survived our inclusion/exclusion criteria – including the validation analysis – consisted of entirely male subjects, most of which were younger adults.

Another limitation to our analysis is the inclusion of samples from a variety of cortical tissues. Four of the included studies used a dissection labeled simply *“cerebral cortex”*, whereas four studies used regional dissections (three frontal cortex, one anterior cingulate). Cortical regions vary in cell type composition and function and are likely to show divergent responses to SD – for example, in older teens and adults the frontal cortex shows the largest build up in SWA in response to homeostatic sleep drive (Jenni and Carskadon, 2007). However, due to the nature of meta-analysis, our results emphasize differential expression that is common across these tissues, and we did not observe obvious clustering related to tissue type in the differential expression of our DEGs. By using a *“cerebral cortex”* dataset as validation, differential expression that was driven by the frontal cortical data was likely excluded from the final “validated” DEG list. Moreover, as our meta-analysis focused on bulk dissections, differential expression specific to any particular cell type or cortical layer is also likely to have been diluted or obscured (*e.g.,* (Ford et al., 2023; Kim et al., 2022; Nakata et al., 2024; Vanrobaeys et al., 2023)). As more data from regional or cell-type specific methods becomes available, future meta-analyses may shed better light on these topics.

Finally, the *Brain Data Alchemy* meta-analysis pipeline was designed as a stopgap measure to address widespread issues with false positive and false negative results in transcriptional profiling studies due to the common use of small sample sizes. To accelerate the generation of higher confidence differential expression results, the pipeline makes transcriptional profiling meta-analysis more efficient and accessible to researchers with limited computational skills (Hagenauer et al., 2024a). Due to access to a large validation dataset, our current study was able to reaffirm the utility and validity of this pipeline. However, the *Brain Data Alchemy* pipeline does introduce several notable limitations. First, the pipeline piggybacks on the curation, re-annotation, and re-analysis efforts of the *Gemma* project, and thus is dependent on the representation of datasets in the *Gemma* database. Although this database is large (>19,000 datasets), it does not encompass all public brain transcriptional profiling datasets, as demonstrated by the absence of the *GSE114845* dataset used as validation. Second, as the pipeline draws from both microarray and RNA-Seq data, it emphasizes gene-level summarized expression from protein coding genes, neglecting alternative splicing and non-coding transcripts (Ford et al., 2023; Wang et al., 2010). This dependency is also likely to bias results away from genes with the lowest levels of expression that are better detected using methods such as qPCR (Medina et al., 2022). That said, by drawing from a large sample size, we can cut through the noise present in measurements from low-level expressed genes, with a few of our validated DEGs having average expression levels less than one count per million (**Tables 2&3**, **Figure S4**). Finally, as with all transcriptional profiling studies, it is important to note that transcript levels may not necessarily reflect protein levels or protein activity, especially since SD is known to affect translation and protein synthesis (Havekes et al., 2012; Lyons et al., 2023).

## Conclusion

In conclusion, by performing a meta-analysis of eight publicly available transcriptional profiling studies characterizing the effects of acute SD on the murine cerebral cortex (collective *n*=293) and validating the results, we improved the reliability and generalizability of our inferences regarding SD effects on cortical gene expression. By releasing the full meta-analysis results (**Table S1:** 16,255 genes, with 182 DEGs) and gene set enrichment results (**Table S2:** 10,436 gene sets, with 236 with FDR<0.05), we provide a useful resource for researchers interested in understanding the molecular correlates of enforced wakefulness and its associated effects on homeostatic sleep drive, mood, cognition, stress response, vascular and immune function. This database may further serve as a useful comparison for cortical transcriptional profiling results from patients with conditions characterized by chronic sleep disruption, including psychiatric, neurodegenerative, and neurodevelopmental disorders. That said, it is important to note that the chronic SD exposure associated with chronic illness is more likely than acute SD exposure to overload the allostatic processes that promote stability in response to challenge, including stress, metabolic, and immune systems (McEwen and Karatsoreos, 2022), driving cerebrovascular dysfunction and altered brain morphology (Kreutzmann et al., 2015; Mahalakshmi et al., 2020). Future directions include elucidating mechanisms by which identified DEGs might mediate behavioral and physiological consequences of SD, and exploring translation from animal models to clinical applications.

## Acknowledgements

We would like to thank our reviewers for providing us with valuable feedback that inspired many of our follow-up analyses, including the validation analysis, and guided our interpretation of the results. This work was completed as part of the *Brain Data Alchemy Project* and supported by the Hope for Depression Research Foundation (HDRF: HA), Grinnell College Center for Careers, Life, and Service (CAR, JX, EH, DMN), the International Brain Research Organization (IBRO) and Faculty for Undergraduate Neuroscience (FUN) (DMN), National Institute on Drug Abuse (NIDA U01 DA043098: HA), and the Pritzker Neuropsychiatric Disorders Research Foundation (HA & SJW). Funders and sponsors had no active role in the review.

## CRediT Statement

CAR – Conceptualization, Methodology, Software, Formal analysis, Investigation, Data Curation, Writing – Original Draft, Writing – Review & Editing, Visualization MHH: Conceptualization, Methodology, Software, Formal analysis, Investigation, Writing – Original Draft, Writing – Review & Editing, Visualization, Validation, Supervision, Project administration

JX: Methodology, Writing – Review & Editing

EH: Methodology, Writing – Review & Editing

DMH: Methodology, Writing – Review & Editing

AS: Methodology, Writing – Review & Editing

EIF: Methodology, Writing – Review & Editing

SJW: Writing – Review & Editing, Funding acquisition

HA: Writing – Review & Editing, Supervision, Funding acquisition

## Data Availability Statement

All datasets used in this publication are publicly available in the National Center for Biotechnology Information (NCBI) Repository Gene Expression Omnibus (GEO, https://www.ncbi.nlm.nih.gov/geo/) under the accession numbers GSE6514, GSE33491, GSE78215, GSE93041, GSE113754, GSE128770, GSE132076, GSE144957, and GSE114845.

