## Supplement for "A Meta-Analysis of the Effects of Acute Sleep Deprivation on the Cortical Transcriptome in Rodent Models"

*Shared first authorship

Direct correspondence to:

Megan Hagenauer, Ph.D.
Michigan Neuroscience Institute
BSRB
109 Zina Pitcher Pl
University of Michigan
Ann Arbor, MI 48109 USA
**

### Supplementary Methods

#### *Data Processing Procedures within Gemma:*

***Data Preprocessing:*** Data preprocessing by *Gemma* includes up-to-date microarray probe alignment and RNA-Seq read mapping (Lim et al., 2021a). For RNA-Seq FASTQ files, read adapters are trimmed with Cutadapt (Martin, 2011) and reads aligned with STAR (Dobin et al., 2013) and quantified using RSEM (Li and Dewey, 2011). For microarray data, different probe-to-gene mapping strategies are applied depending on platform. For Affymetrix datasets such as those referenced in our paper, data is extracted from the raw .CEL files using the RMA algorithm (Irizarry et al., 2003) within Affymetrix Analysis Power Tools. Probe sets are collapsed to a single representative sequence, resolving overlaps (Barnes et al., 2005), and aligned to the reference genome with BLAT (Kent, 2002). Alignments are then filtered for specificity, and mapped to transcripts using UCSC Golden Path (Lim et al., 2021a).

***Quality Control:*** For gene-level quality control, the *Gemma* pipeline screens out genes lacking variance in expression levels (<70% distinct values), but otherwise does not apply filtering based on expression level. This inclusion of data from low level expressed genes can increase the risk for false positive and false negative findings in differential expression analyses due to floor effects in transcriptional profiling data, but filtering out low expression data poses its own risks in biasing inferences – in particular, the expression for many neurotransmission-related genes is very low in brain tissue, despite being of particular import for understanding neuropsychiatric conditions (Medina, Hagenauer, Krolewski et al. 2023).

Sample-level quality control is conducted in *Gemma* using a procedure that can be applied similarly across platforms. Samples are considered potential outliers if their adjusted median sample-sample correlation is outside the interquartile range of the sample-sample correlations for the sample with the closest median correlation to them (Lim et al., 2021b). Processing batches are defined automatically using information in the raw data files (*microarrays:* clusters of date stamps, *RNA-Seq:* FASTQ header information such as "device", "run", "flow cell" and/or "lane"), or obtained manually from associated supplementary files or publications, and screened for correlation with the top three principal components of variation in the expression data (Lim et al., 2021b). Batch correction is conducted using an in-house implementation of the ComBat algorithm (Johnson et al., 2007). Batch correction is not performed when no substantial batch effect is detected or when batches are confounded with experimental design factors, in which case the dataset is flagged or split.

### Supplemental Tables

***Table S1. The full meta-analysis results (16,290 genes, 16,255 stable meta-analysis estimates).*** *This .xlsx file includes two worksheets: 1) The worksheet "SleepDeprivation_MetaAnalysisResults" provides the full meta-analysis results, with each row representing the results for one gene, and each column providing either gene annotation or meta-analysis statistical output. The results are ordered by p-value, so that the top rows in the worksheet are the genes with the smallest p-values. 2) The worksheet "ColumnDefinitions"* *provides the definitions for the variables present in each column in "SleepDeprivation_MetaAnalysisResults".*

***Table S2. Full differential expression results from the re-analysis of GSE114845 (19,798 genes).*** *This .xlsx file includes two worksheets: 1) The worksheet "SleepDep_GSE114845_DEResults" provides the full differential expression results from our re-analysis of GSE114845, with each row representing the results for one gene, and each column providing either gene annotation or differential expression statistical output. The results are ordered by p-value, so that the top rows in the worksheet are the genes with the smallest p-values. 2) The worksheet "ColumnDefinitions" provides the definitions for the variables present in each column in "SleepDep_GSE114845_DEResults".*

| | Meta-Analysis: SD | | | | | | | | Validation: GSE114845 SD | | | | | | Exploratory |
|---|---|---|---|---|---|---|---|---|---|---|---|---|---|---|---|
| **Gene Symbol** | **#** | **Log2FC** | **CI_lb** | **CI_ub** | **SE** | **Tstat** | **Pval** | **FDR** | **Expr** | **Log2FC** | **SE** | **Tstat** | **Pval** | **FDR** | **RS Log2FC** |
| ***Rimoc1*** | 16 | **-0.081** | -0.104 | -0.059 | 0.011 | -7.38 | 1.33E-12 | 1.96E-08 | NA | NA | NA | NA | NA | NA | 0.017 |
| ***Tmod3*** | 18 | **-0.111** | -0.151 | -0.070 | 0.021 | -5.37 | 8.00E-08 | 1.00E-04 | 4.97 | **0.096** | 0.021 | 4.57 | 1.60E-05 | 4.18E-05 | 0.080 |
| ***Lrrc75a*** | 18 | **-0.116** | -0.159 | -0.073 | 0.022 | -5.28 | 1.28E-07 | 1.15E-04 | 3.90 | **0.077** | 0.030 | 2.60 | 1.08E-02 | 1.88E-02 | 0.078 |
| ***Mxi1*** | 18 | **-0.061** | -0.085 | -0.037 | 0.012 | -5.00 | 5.81E-07 | 3.79E-04 | 6.20 | **0.040** | 0.016 | 2.50 | 1.42E-02 | 2.41E-02 | -0.012 |
| ***Khnyn*** | 18 | **-0.091** | -0.130 | -0.052 | 0.020 | -4.55 | 5.38E-06 | 2.31E-03 | 4.96 | 0.009 | 0.039 | 0.22 | 8.23E-01 | 8.65E-01 | 0.061 |
| ***C130071C03Rik*** | 18 | **-0.126** | -0.181 | -0.070 | 0.028 | -4.44 | 9.20E-06 | 3.48E-03 | 6.20 | ***0.052*** | 0.025 | 2.08 | 4.02E-02 | 6.26E-02 | ***0.114*** |
| ***Pdk4*** | 18 | **-0.171** | -0.248 | -0.095 | 0.039 | -4.39 | 1.12E-05 | 3.89E-03 | 3.47 | **0.178** | 0.047 | 3.80 | 2.69E-04 | 5.95E-04 | -0.003 |
| ***Nrg2*** | 18 | **-0.084** | -0.123 | -0.046 | 0.020 | -4.30 | 1.69E-05 | 5.20E-03 | 2.72 | 0.067 | 0.032 | 2.10 | 3.83E-02 | 6.01E-02 | 0.072 |
| ***Rufy3*** | 18 | **-0.049** | -0.072 | -0.026 | 0.012 | -4.18 | 2.86E-05 | 8.17E-03 | 7.75 | 0.018 | 0.010 | 1.88 | 6.28E-02 | 9.42E-02 | 0.010 |
| ***Pcdhb3*** | 18 | **-0.109** | -0.161 | -0.058 | 0.026 | -4.18 | 2.91E-05 | 8.17E-03 | 3.06 | 0.026 | 0.063 | 0.42 | 6.75E-01 | 7.41E-01 | -0.017 |
| ***Pqlc3*** | 18 | **-0.118** | -0.175 | -0.061 | 0.029 | -4.04 | 5.26E-05 | 1.09E-02 | 2.33 | **0.227** | 0.079 | 2.87 | 5.12E-03 | 9.37E-03 | 0.097 |
| ***Rassf4*** | 18 | **-0.099** | -0.147 | -0.051 | 0.025 | -4.03 | 5.68E-05 | 1.13E-02 | 3.43 | 0.060 | 0.045 | 1.35 | 1.79E-01 | 2.41E-01 | 0.034 |
| ***Eml1*** | 18 | **-0.062** | -0.093 | -0.031 | 0.016 | -3.96 | 7.47E-05 | 1.36E-02 | 5.97 | 0.009 | 0.015 | 0.59 | 5.55E-01 | 6.31E-01 | 0.050 |
| ***Fam13c*** | 18 | **-0.081** | -0.121 | -0.040 | 0.020 | -3.94 | 8.30E-05 | 1.47E-02 | 5.98 | **0.110** | 0.022 | 5.09 | 2.04E-06 | 5.88E-06 | 0.051 |
| ***Slain1*** | 16 | **-0.083** | -0.125 | -0.042 | 0.021 | -3.93 | 8.48E-05 | 1.47E-02 | 5.17 | -0.039 | 0.022 | -1.79 | 7.70E-02 | 1.13E-01 | -0.022 |
| ***Sc5d*** | 18 | **-0.090** | -0.135 | -0.045 | 0.023 | -3.91 | 9.05E-05 | 1.52E-02 | 5.75 | 0.054 | 0.055 | 0.98 | 3.30E-01 | 4.08E-01 | 0.052 |
| ***Acadm*** | 18 | **-0.081** | -0.122 | -0.040 | 0.021 | -3.90 | 9.50E-05 | 1.56E-02 | 4.71 | -0.006 | 0.014 | -0.42 | 6.77E-01 | 7.42E-01 | 0.042 |
| ***Rufy2*** | 18 | **-0.055** | -0.083 | -0.027 | 0.014 | -3.85 | 1.17E-04 | 1.75E-02 | 6.79 | -0.035 | 0.018 | -1.91 | 5.92E-02 | 8.93E-02 | 0.018 |
| ***Septin8*** | 18 | **-0.056** | -0.084 | -0.027 | 0.014 | -3.85 | 1.20E-04 | 1.77E-02 | 8.56 | **0.074** | 0.014 | 5.11 | 1.89E-06 | 5.49E-06 | 0.049 |
| ***Ccdc30*** | 18 | **-0.067** | -0.101 | -0.032 | 0.018 | -3.78 | 1.57E-04 | 2.09E-02 | 4.13 | **0.044** | 0.018 | 2.46 | 1.58E-02 | 2.66E-02 | 0.047 |
| ***Mogs*** | 18 | **-0.081** | -0.124 | -0.039 | 0.022 | -3.76 | 1.69E-04 | 2.20E-02 | 4.43 | **0.055** | 0.019 | 2.86 | 5.28E-03 | 9.65E-03 | 0.069 |
| ***Car2*** | 18 | **-0.112** | -0.171 | -0.054 | 0.030 | -3.75 | 1.74E-04 | 2.23E-02 | 7.24 | -0.091 | 0.048 | -1.89 | 6.22E-02 | 9.33E-02 | -0.032 |
| ***Bean1*** | 18 | **-0.110** | -0.168 | -0.052 | 0.029 | -3.74 | 1.83E-04 | 2.29E-02 | 3.95 | -0.001 | 0.026 | -0.04 | 9.72E-01 | 9.80E-01 | 0.039 |
| ***Tal1*** | 18 | **-0.141** | -0.214 | -0.067 | 0.038 | -3.74 | 1.86E-04 | 2.29E-02 | 0.86 | 0.068 | 0.051 | 1.33 | 1.88E-01 | 2.51E-01 | 0.144 |
| ***Rabl2*** | 18 | **-0.074** | -0.113 | -0.035 | 0.020 | -3.72 | 1.96E-04 | 2.38E-02 | 4.79 | 0.032 | 0.025 | 1.32 | 1.89E-01 | 2.53E-01 | 0.086 |
| ***Cep120*** | 18 | **-0.042** | -0.065 | -0.020 | 0.011 | -3.69 | 2.20E-04 | 2.65E-02 | 5.70 | 0.017 | 0.013 | 1.24 | 2.17E-01 | 2.85E-01 | -0.030 |
| ***Nol4l*** | 18 | **-0.085** | -0.131 | -0.039 | 0.023 | -3.65 | 2.61E-04 | 2.97E-02 | 5.85 | **0.092** | 0.023 | 4.05 | 1.10E-04 | 2.57E-04 | 0.083 |
| ***Pnrc1*** | 18 | **-0.155** | -0.238 | -0.071 | 0.042 | -3.64 | 2.72E-04 | 3.03E-02 | 4.47 | 0.005 | 0.024 | 0.19 | 8.50E-01 | 8.86E-01 | 0.068 |
| ***BC002059*** | 18 | **-0.158** | -0.245 | -0.072 | 0.044 | -3.59 | 3.35E-04 | 3.59E-02 | 2.70 | **0.067** | 0.025 | 2.71 | 8.08E-03 | 1.43E-02 | -0.023 |
| ***Zfp658*** | 18 | **-0.185** | -0.286 | -0.083 | 0.052 | -3.57 | 3.55E-04 | 3.73E-02 | 3.11 | **0.341** | 0.106 | 3.23 | 1.76E-03 | 3.46E-03 | 0.169 |
| ***Srpk1*** | 18 | **-0.049** | -0.077 | -0.022 | 0.014 | -3.54 | 3.94E-04 | 4.01E-02 | 6.56 | 0.000 | 0.015 | -0.03 | 9.77E-01 | 9.83E-01 | 0.061 |
| ***Efnb2*** | 18 | **-0.049** | -0.076 | -0.022 | 0.014 | -3.52 | 4.30E-04 | 4.20E-02 | 5.75 | ***-0.038*** | 0.018 | -2.10 | 3.84E-02 | 6.02E-02 | -0.013 |
| ***2410021H03Rik*** | 16 | **-0.108** | -0.168 | -0.048 | 0.031 | -3.52 | 4.38E-04 | 4.25E-02 | -0.15 | 0.090 | 0.092 | 0.97 | 3.35E-01 | 4.14E-01 | 0.194 |
| ***Sncaip*** | 18 | **-0.100** | -0.156 | -0.043 | 0.029 | -3.48 | 5.10E-04 | 4.75E-02 | 1.72 | -0.049 | 0.043 | -1.12 | 2.67E-01 | 3.41E-01 | 0.100 |
| ***Mtmr14*** | 18 | **-0.065** | -0.102 | -0.028 | 0.019 | -3.46 | 5.32E-04 | 4.90E-02 | 3.69 | -0.030 | 0.025 | -1.18 | 2.43E-01 | 3.14E-01 | 0.036 |
| ***Zfp354a*** | 18 | **-0.089** | -0.140 | -0.039 | 0.026 | -3.46 | 5.50E-04 | 4.95E-02 | 3.00 | -0.030 | 0.028 | -1.07 | 2.89E-01 | 3.65E-01 | -0.026 |
| ***Dtx4*** | 18 | **-0.081** | -0.126 | -0.035 | 0.023 | -3.45 | 5.58E-04 | 4.99E-02 | 5.98 | **0.070** | 0.017 | 4.08 | 9.87E-05 | 2.32E-04 | 0.075 |

***Table S3. Genes that were downregulated in the murine cortex following SD in our meta-analysis of public transcriptional profiling data (FDR<0.05, collective n=293) but not validated by a separate large RNA-Seq study (GSE114845, n=86 samples representing 222 mice).*** *Differential expression was considered to be not validated if 1) the opposite direction of effect was observed in GSE114845 than what was observed in the meta-analysis or 2) the same direction of effect was observed in GSE114845 as in the meta-analysis, but the differential expression was not significant (FDR<0.05). Similar to* ***Table 2****, within the Log(2)Fold Change (Log2FC) columns, blue is used to highlight down-regulation, pink is used to highlight up-*

*regulation, bold text indicates FDR<0.05, and bold/italic text indicates nominal significance (p<0.05). Column definitions: #= Number of differential expression results (SD vs. control statistical contrasts) that contributed to the meta-analysis estimate for that gene; Log2FC=estimated SD vs. control Log2 Fold Change; CI_lb & CI_ub=Lower and upper bound for the 95% confidence interval for the Log2FC; SE=Standard error for the Log2FC, Tstat=T-statistic for the Log2FC, Pval=Nominal p-value for the Log2FC, FDR=False discovery rate for the Log2FC (q-value or adjusted p-value), Expr=Average Log2 expression (counts per million: cpm) for the gene, RS Log2FC: estimated moderating effect (Log(2) Fold Change) of recovery sleep (RS) on the effect of SD within our exploratory meta-analysis.*

| | Meta-Analysis: SD | | | | | | | | Validation: GSE114845 SD | | | | | | Exploratory |
|---|---|---|---|---|---|---|---|---|---|---|---|---|---|---|---|
| Gene Symbol | # | Log2FC | CI_lb | CI_ub | SE | Tstat | Pval | FDR | Expr | Log2FC | SE | Tstat | Pval | FDR | RS Log2FC |
| ***Thg1l*** | 18 | **0.103** | 0.074 | 0.132 | 0.015 | 7.01 | 2.41E-12 | 1.96E-08 | 3.90 | 0.052 | 0.041 | 1.27 | 2.07E-01 | 2.73E-01 | -0.064 |
| ***Fancf*** | 18 | **0.109** | 0.071 | 0.147 | 0.019 | 5.61 | 2.07E-08 | 6.75E-05 | 0.95 | **-0.249** | 0.050 | -4.95 | 3.57E-06 | 1.00E-05 | -0.043 |
| ***Gm14285*** | 18 | **0.182** | 0.116 | 0.248 | 0.034 | 5.39 | 6.95E-08 | 1.00E-04 | -0.78 | 0.088 | 0.136 | 0.64 | 5.21E-01 | 6.00E-01 | -0.062 |
| ***Exosc4*** | 18 | **0.146** | 0.093 | 0.200 | 0.027 | 5.37 | 7.73E-08 | 1.00E-04 | 3.57 | **-0.062** | 0.025 | -2.51 | 1.38E-02 | 2.35E-02 | -0.026 |
| ***4931429L15Rik*** | 18 | **0.136** | 0.086 | 0.186 | 0.026 | 5.31 | 1.09E-07 | 1.11E-04 | NA | NA | NA | NA | NA | NA | -0.014 |
| ***Card6*** | 18 | **0.145** | 0.080 | 0.210 | 0.033 | 4.36 | 1.31E-05 | 4.37E-03 | 1.13 | -0.042 | 0.040 | -1.05 | 2.95E-01 | 3.71E-01 | ***-0.160*** |
| ***Tmem81*** | 18 | **0.132** | 0.072 | 0.191 | 0.030 | 4.33 | 1.51E-05 | 4.84E-03 | 1.59 | **-0.142** | 0.051 | -2.77 | 6.90E-03 | 1.24E-02 | -0.063 |
| ***Cspg4*** | 18 | **0.117** | 0.062 | 0.173 | 0.028 | 4.16 | 3.17E-05 | 8.62E-03 | 4.55 | 0.022 | 0.031 | 0.72 | 4.72E-01 | 5.52E-01 | -0.112 |
| ***Jund*** | 18 | **0.083** | 0.044 | 0.122 | 0.020 | 4.15 | 3.34E-05 | 8.78E-03 | 6.44 | 0.043 | 0.037 | 1.18 | 2.43E-01 | 3.14E-01 | -0.047 |
| ***Bphl*** | 18 | **0.060** | 0.032 | 0.089 | 0.015 | 4.13 | 3.56E-05 | 9.07E-03 | 3.88 | **-0.077** | 0.024 | -3.24 | 1.68E-03 | 3.31E-03 | 0.000 |
| ***Lsm3*** | 18 | **0.110** | 0.057 | 0.162 | 0.027 | 4.10 | 4.16E-05 | 9.98E-03 | 3.21 | 0.048 | 0.027 | 1.80 | 7.46E-02 | 1.10E-01 | -0.103 |
| ***Gemin6*** | 18 | **0.122** | 0.064 | 0.181 | 0.030 | 4.09 | 4.33E-05 | 1.00E-02 | 1.66 | -0.040 | 0.033 | -1.22 | 2.25E-01 | 2.93E-01 | -0.085 |
| ***Sh3bp2*** | 18 | **0.110** | 0.057 | 0.163 | 0.027 | 4.05 | 5.21E-05 | 1.09E-02 | 1.44 | 0.065 | 0.049 | 1.34 | 1.83E-01 | 2.45E-01 | -0.011 |
| ***Psmc2*** | 18 | **0.052** | 0.026 | 0.077 | 0.013 | 3.99 | 6.59E-05 | 1.23E-02 | 6.23 | -0.016 | 0.019 | -0.86 | 3.90E-01 | 4.71E-01 | -0.034 |
| ***Maip1*** | 18 | **0.084** | 0.042 | 0.125 | 0.021 | 3.95 | 7.95E-05 | 1.42E-02 | 4.16 | 0.014 | 0.018 | 0.79 | 4.32E-01 | 5.13E-01 | -0.014 |
| ***Znhit3*** | 18 | **0.081** | 0.041 | 0.122 | 0.021 | 3.91 | 9.05E-05 | 1.52E-02 | 3.71 | 0.025 | 0.022 | 1.14 | 2.59E-01 | 3.32E-01 | 0.008 |
| ***Rhob*** | 18 | **0.107** | 0.052 | 0.162 | 0.028 | 3.84 | 1.21E-04 | 1.77E-02 | 7.28 | 0.025 | 0.024 | 1.05 | 2.97E-01 | 3.73E-01 | ***-0.140*** |
| ***Clec18a*** | 18 | **0.129** | 0.063 | 0.195 | 0.034 | 3.81 | 1.39E-04 | 1.89E-02 | 3.29 | 0.064 | 0.064 | 1.00 | 3.22E-01 | 4.00E-01 | 0.012 |
| ***Gstm6*** | 18 | **0.126** | 0.060 | 0.191 | 0.034 | 3.75 | 1.76E-04 | 2.24E-02 | 1.04 | 0.103 | 0.060 | 1.71 | 9.14E-02 | 1.32E-01 | -0.030 |
| ***Nsmce3*** | 18 | **0.075** | 0.036 | 0.114 | 0.020 | 3.74 | 1.82E-04 | 2.29E-02 | 4.19 | -0.028 | 0.022 | -1.24 | 2.19E-01 | 2.86E-01 | ***-0.096*** |
| ***Gm6225*** | 16 | **0.276** | 0.131 | 0.420 | 0.074 | 3.74 | 1.87E-04 | 2.29E-02 | 1.92 | -0.016 | 0.038 | -0.41 | 6.85E-01 | 7.48E-01 | -0.241 |
| ***Tprn*** | 18 | **0.095** | 0.043 | 0.146 | 0.026 | 3.61 | 3.11E-04 | 3.42E-02 | 3.58 | 0.007 | 0.030 | 0.24 | 8.10E-01 | 8.55E-01 | -0.115 |
| ***Hrob*** | 18 | **0.103** | 0.047 | 0.158 | 0.029 | 3.59 | 3.26E-04 | 3.54E-02 | 0.28 | 0.025 | 0.071 | 0.35 | 7.29E-01 | 7.86E-01 | -0.082 |
| ***Kcng2*** | 18 | **0.172** | 0.077 | 0.267 | 0.049 | 3.55 | 3.82E-04 | 3.96E-02 | 1.47 | **-0.224** | 0.058 | -3.87 | 2.06E-04 | 4.62E-04 | -0.073 |
| ***Gjc1*** | 18 | **0.141** | 0.063 | 0.220 | 0.040 | 3.54 | 4.07E-04 | 4.06E-02 | 2.21 | 0.062 | 0.036 | 1.73 | 8.72E-02 | 1.27E-01 | ***-0.202*** |
| ***Smg8*** | 18 | **0.055** | 0.024 | 0.087 | 0.016 | 3.50 | 4.62E-04 | 4.43E-02 | 3.77 | 0.039 | 0.022 | 1.79 | 7.74E-02 | 1.14E-01 | 0.016 |
| ***Rps20*** | 18 | **0.082** | 0.036 | 0.128 | 0.023 | 3.50 | 4.71E-04 | 4.46E-02 | 5.96 | **-0.097** | 0.028 | -3.47 | 7.97E-04 | 1.65E-03 | -0.071 |
| ***AI429214*** | 16 | **0.116** | 0.051 | 0.182 | 0.033 | 3.48 | 4.96E-04 | 4.67E-02 | 1.79 | 0.073 | 0.078 | 0.93 | 3.53E-01 | 4.33E-01 | -0.102 |
| ***Rhou*** | 18 | **0.126** | 0.055 | 0.198 | 0.036 | 3.48 | 5.00E-04 | 4.68E-02 | 5.21 | -0.015 | 0.029 | -0.51 | 6.10E-01 | 6.82E-01 | -0.121 |
| ***BC003965*** | 18 | **0.065** | 0.028 | 0.103 | 0.019 | 3.46 | 5.48E-04 | 4.95E-02 | 4.67 | 0.014 | 0.021 | 0.66 | 5.10E-01 | 5.89E-01 | -0.041 |

***Table S4. Genes that were upregulated in the frontal cortex following sleep deprivation in our meta-analysis of public transcriptional profiling data (FDR<0.05, collective n=293) but not validated by a separate large RNA-Seq study (GSE114845, n=86 samples representing 222 mice).*** *Differential expression was considered to be not validated if 1) the opposite direction of effect was observed in GSE114845 than what was observed in the meta-analysis or 2) the same direction of effect was observed in GSE114845 as in the meta-analysis, but the differential expression was not significant (FDR<0.05). Column definitions are the same as* ***Table S3.***

***Table S5: The full fast Gene Set Enrichment Analysis (fGSEA) results (10,436 gene sets).*** *This .xlsx file includes three worksheets: 1) The worksheet "fGSEA_Results_Directional" provides the full fGSEA results for the directional analysis, with each row representing the results for one gene set, and each column providing the fGSEA statistical output. The results are ordered by p-value, so that the top rows in the worksheet are the gene sets with the smallest p-values. 2) The* *worksheet "fGSEA_Results_Nondirectional" provides the full fGSEA results for the non-directional analysis in the same format as the directional analysis output. 3) The worksheet "ColumnDefinitions" provides the definitions for the variables present in each column in "fGSEA_Results_Directional" and "fGSEA_Results_NonDirectional".*

***Table S6. The full exploratory meta-analysis results (16,290 genes, 16,248 stable meta-analysis estimates).*** *This .xlsx file includes two worksheets: 1) The worksheet "ExploratoryMeta_FullResults" provides the full exploratory meta-analysis results, with each row representing the results for one gene, and each column providing either gene annotation or meta-analysis statistical output. The results are ordered by p-value for the effect of recovery sleep (RS), so that the top rows in the worksheet are the genes with the smallest p-values. 2) The worksheet "ColumnDefinitions" provides the definitions for the variables present in each column in "ExploratoryMeta_FullResults".*

| | Current Meta-Analysis | GSE114845 Validation | Juszczak & Stankiewicz 2018 | Jaszczyk et al., 2023 | | | Jaszczyk et al., 2025 | Meta-Analysis: Stankiewicz et al.., 2022 | | | |
|---|---|---|---|---|---|---|---|---|---|---|---|
| | Sleep Deprivation | | Responsive to Glucocorticoids | Acute 12h Corticosterone | | | Chronic Daily Corticosterone | Responsive to Stress | | | |
| Gene | Log2FC | Validated? | Review of Literature | 1 h post | 5 h post | 9 h post | Pooled: 5 days, 14 days, or 28 days | Overall Stress-responsive | Acute Stress-responsive | Medium Stress-responsive | Prolonged Stress-responsive |
| ***Nr3c1*** | -0.122 | Down | Down | Down | Down | Down | Down | | | | |
| ***Car2*** | -0.112 | No | | Down | Down | | Down | | | | Down |
| ***Lrrc75a*** | -0.116 | No | Down | Down | Down | | Down | | | | |
| ***Fa2h*** | -0.106 | Down | | Down | Down | | Down | Up&Down | | | Up&Down |
| ***Septin8*** | -0.056 | No | | Down | Down | Down | | | | | |
| ***Herpud2*** | -0.051 | Down | | Down | Down | Down | | | | | |
| ***Cep120*** | -0.042 | No | | Down | Down | Down | | Up&Down | | Up&Down | |
| ***Zfp521*** | -0.108 | Down | | Down | Down | | Down | | | | |
| ***Sox21*** | -0.263 | Down | | Down | Down | | | Up&Down | | | Up&Down |
| ***Tbc1d31*** | -0.111 | Down | | Down | Down | | | | | | |
| ***Smc2*** | -0.098 | Down | | Down | Down | | | | | | |
| ***Il17rb*** | -0.093 | Down | | Down | | Down | | | | | |
| ***Rufy2*** | -0.055 | No | | Down | Up&Down | Down | | | | | |
| ***Ahcy*** | -0.095 | Down | | | Down | Down | | | | | |
| ***Rbm5*** | -0.073 | Down | | | Down | Down | | | | | |
| ***Mpzl1*** | -0.112 | Down | | Down | | | Down | | | | |
| ***Trim59*** | -0.169 | Down | | Down | | | Down | | | | |
| ***2810459M11Rik*** | -0.157 | Down | | Down | | | Down | | | | |
| ***Ablim3*** | -0.148 | Down | | | | Down | | | | | Up&Down |
| ***Vps26c*** | -0.081 | Down | | | | | | Up&Down | Down | | |
| ***Mrtfa*** | 0.102 | Up | | Up | | Down | Up | | | | |
| ***Card6*** | 0.145 | No | | Up | Up | Down | | | | | |
| ***Ramp2*** | 0.236 | Up | | Up | Up | Down | | | | | |
| ***Sorcs3*** | 0.147 | Up | | | | Up | | | | Up&Down | |
| ***Mchr1*** | 0.095 | Up | | Up | | | Up | | | | |
| ***Gata2*** | 0.135 | Up | | Up | | | Up | | | | |
| ***Vip*** | 0.133 | Up | | | Up | | Up | Up&Down | | | Up&Down |
| ***Lingo3*** | 0.155 | Up | | Up | | | Up | | | | |
| ***Htr2a*** | 0.158 | Up | | Up | | | Up | | | | |
| ***Mink1*** | 0.043 | Up | | Up | Up | Up | | | | Down | |
| ***Smcr8*** | 0.052 | Up | | Up | | | | | | Up | |
| ***Nt5dc3*** | 0.089 | Up | | Up | Up | | | | | | |
| ***Hrob*** | 0.103 | No | | Up | | Up | | | | | |
| ***Hspa12b*** | 0.153 | Up | | Up | Up | Up&Down | | | | | |
| ***Ppp1r10*** | 0.102 | Up | | Up&Down | Up | Up | | | | | |
| ***Fam110b*** | 0.112 | Up | Up | Up&Down | Up | Up&Down | | | | | |
| ***Tsnax*** | 0.064 | Up | | | Up | Up | | | | | |
| ***Jund*** | 0.083 | No | | | Up | Up | | Up&Down | | | |
| ***Fam78a*** | 0.093 | Up | | | Up | Up | | | | | |
| ***Rhou*** | 0.126 | No | Up | Up | | | Up | | | | |
| ***Ociad2*** | 0.147 | Up | | Up | Up | | Up | | | | |
| ***Arpc2*** | 0.06 | Up | | Up | Up | | Up | | | | |
| ***Pip5k1a*** | 0.043 | Up | | Up | Up | | Up | | | | |
| ***Kcng2*** | 0.172 | No | | Up | Up | | Up | | | | |
| ***Zbtb16*** | 0.255 | Up | | Up | Up | Down | Up | Up&Down | | | Up |
| ***Plcl2*** | 0.057 | Up | | Up | Up | Up | | | | | |
| ***Wnk2*** | 0.065 | Up | | Up | Up | Up | | | | | |
| ***Ddx51*** | 0.084 | Up | | Up | Up | Up | | | | | |
| ***Gstm6*** | 0.126 | No | | Up | | Up | | Up&Down | | Up | |
| ***Prodh*** | 0.123 | Up | | Up | Up | Up | Up | | | | |
| ***Mapk4*** | 0.095 | Up | | Up | Up | Up | Up | | | | |
| ***Rhob*** | 0.107 | No | Up | Up | Up | | Up | | | | |
| ***Gm9925*** | 0.189 | Up | | Up | Up | Up | Up | | | | |
| ***Prr5*** | 0.302 | Up | Up | Up | Up | | Up | | Up&Down | | |
| ***Ntrk2*** | 0.047 | Up | Up&Down | Up | Up | Up | | Up&Down | Up | | Up&Down |
| ***Myo1e*** | 0.106 | Up | | Up | Up | Up | | | | Up | |
| ***Mt2*** | 0.349 | Up | Up | Up | Up | | Up | Up&Down | Up | | |

***Table S7. Many of the genes that are differentially expressed in response to SD are also similarly responsive to glucocorticoids or stress.*** *We compared our list of DEGs for SD to a published database (Table S2 in (Juszczak et al., 2025)) overviewing evidence from two large studies examining the effects of acute (n=48) (Jaszczyk et al., 2023) and chronic daily (n=48)(Juszczak et al., 2025) stress hormone (corticosterone) exposure via drinking water on* *gene expression in the hippocampus, as well as previous evidence from a large vote-counting style meta-analysis of published effects of glucocorticoids (17 studies) (Juszczak and Stankiewicz, 2018) and of published effects of stress (79 animal studies (n=1887) and 3 human studies (n=121)) (Stankiewicz et al., 2022) on brain gene expression as measured using transcriptional profiling. We found that 78 of our DEGs have consistent evidence linking them to either glucocorticoid or stress effects on the brain. Of these, 57 DEGs showed effects of glucocorticoids or stress that were in the same direction as SD (shown above) and another 21 DEGs showed effects that were in the opposing direction from SD (not shown).*

### Supplemental Figures

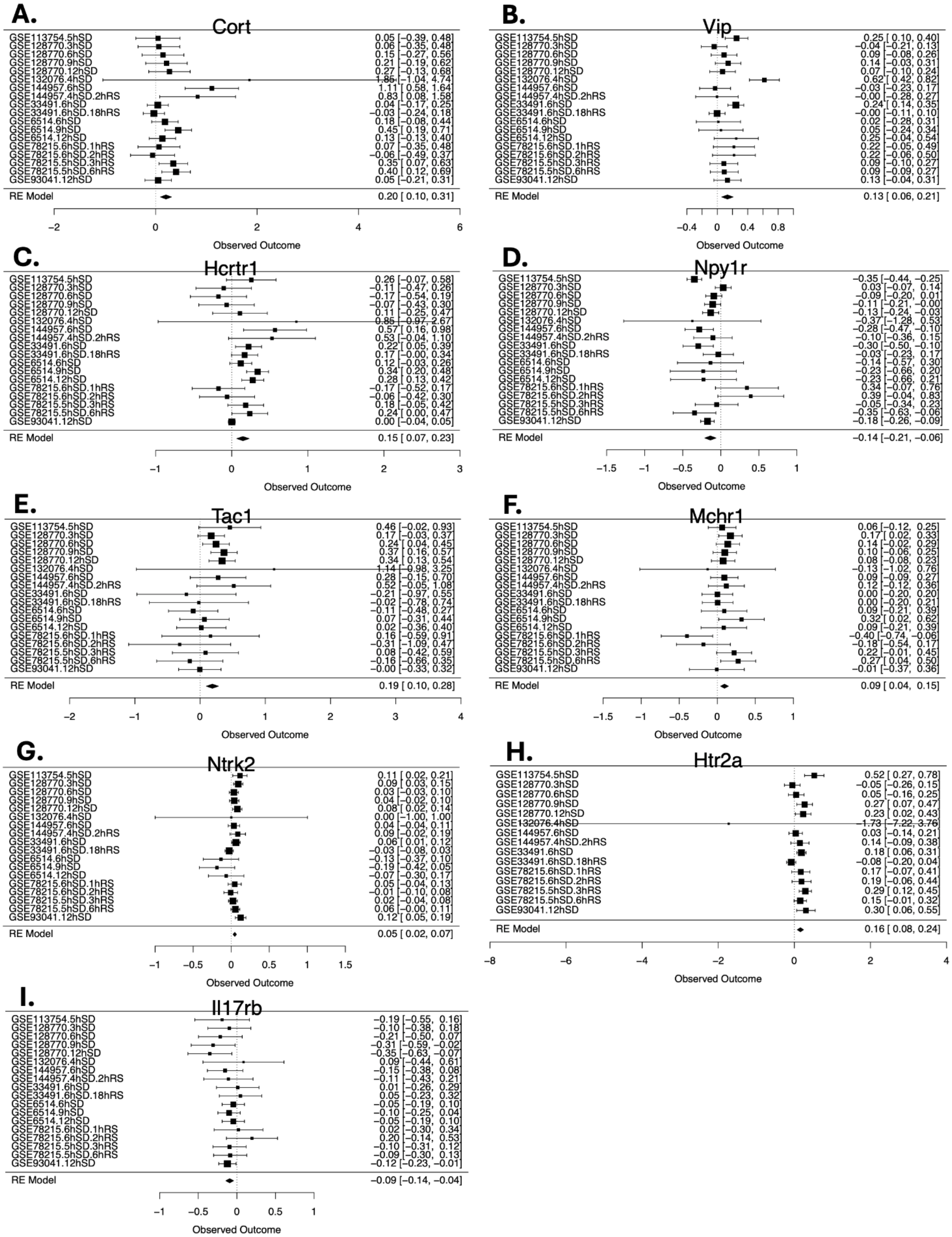

***Figure S1. More example forest plots that show neuropeptide and cell signalling related genes that were consistently differentially expressed in the murine cortex across SD paradigms and***

***experiments in our meta-analysis of public datasets (collective n=293) and validated using an independent large RNA-Seq dataset (GSE114845).*** *Rows illustrate the effect sizes (SD vs. control Log2FC) with squares, with 95% confidence intervals (whiskers), for each of the datasets and the meta-analysis random effects meta-analysis model ("RE Model"). Each study is named using the duration of SD in hours (h), the Gene Expression Omnibus accession number (GSE#), and, if relevant, the duration of recovery sleep (RS) in hours (hr).*

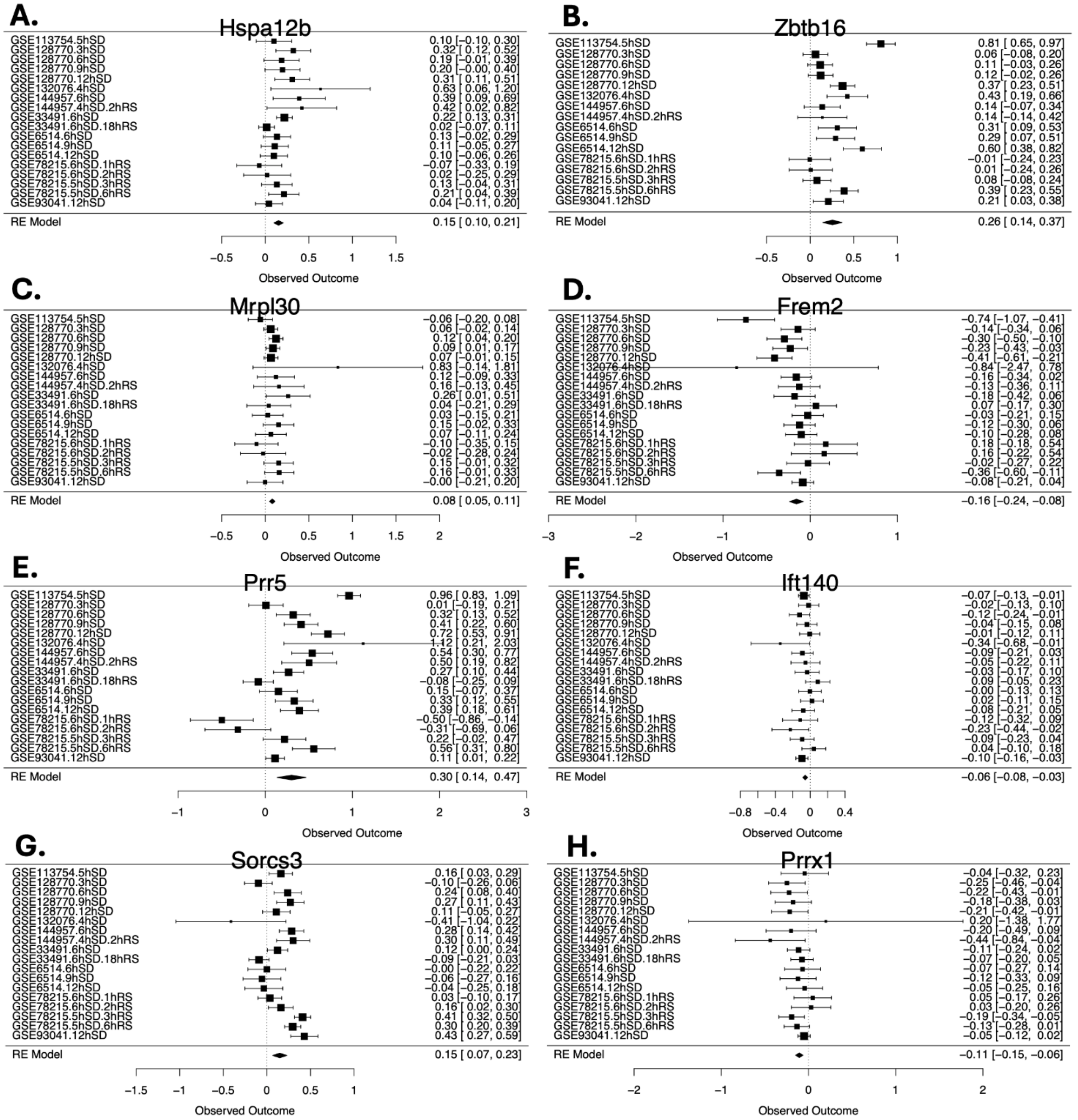


***Figure S2. More example forest plots for other genes mentioned in the main text that were consistently differentially expressed in the murine cortex across SD paradigms and experiments in our meta-analysis of public datasets (collective n=293) and validated using an independent large RNA-Seq dataset (GSE114845).*** *Follows the conventions of* ***Figure S1****.*

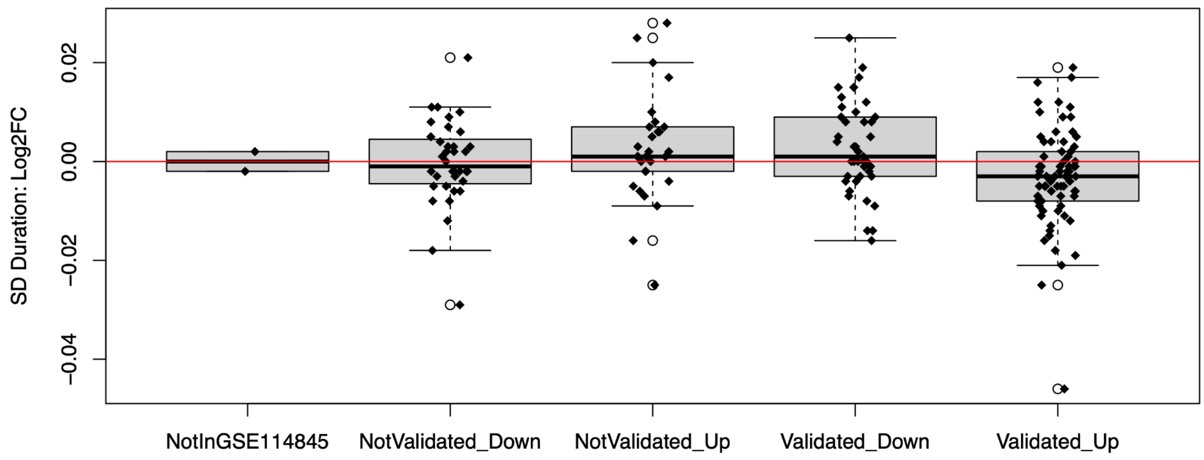


***Figure S3. Exploratory: When examining the effects of sleep deprivation in the exploratory meta-analysis, the 182 genes that showed an effect of SD in the planned meta-analysis (FDR<0.05) did not consistently show an effect of SD Duration (Log2FC: y-axis) in the same direction as the effect of SD in the planned meta-analysis.*** *This was true for both the DEGs that were fully validated using GSE114845 and those that were not.*

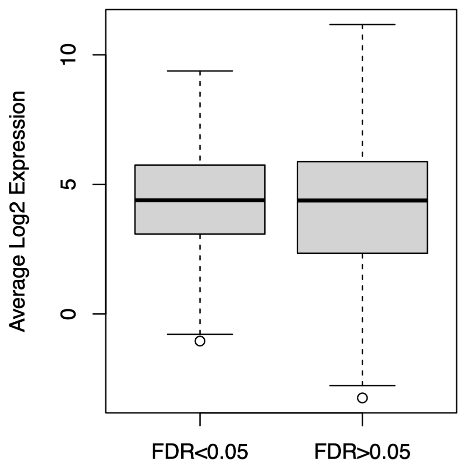


***Figure S4. The genes that are consistently differentially expressed in response to sleep deprivation across datasets in the meta-analysis (FDR<0.05) do not tend to be biased towards genes with high level expression.*** *Referencing the average Log2 expression levels from the large RNA-Seq study that we used as validation for the meta-analysis (GSE114845), we did not find that the genes that were identified as DEGs in our meta-analysis (FDR<0.05) had noticeably higher levels of expression than genes that were not differentially expressed in our meta-analysis, although there may be a slight skew away from the lowest end of the distribution (as would be expected). Likewise, the DEGs that were fully validated using GSE114845 (115 genes)*

*had an average Log2 expression of 4.31+/-0.20 SE, and the DEGs that did not validate using GSE114845 (65 genes) had an average expression of 3.97+/-0.25 SE, showing a slight skew toward higher level expressed genes validating that was not statistically significant.*
